# Confronting spurious evaluations of computational methods in small molecule mass spectrometry

**DOI:** 10.64898/2026.05.03.722532

**Authors:** Vishu Gupta, Chencheng Xu, Ehud Herbst, Fei Wang, David S. Wishart, Michael A. Skinnider

## Abstract

Mass spectrometry-based metabolomics detects thousands of small molecule-associated signals in biological samples, but the vast majority cannot be structurally identified. Mounting interest in this metabolomic “dark matter” has spurred the development of dozens of machine-learning models for structural annotation of small molecules from their MS/MS spectra. Here, we expose a fundamental flaw in the longstanding paradigm by which these models have been evaluated. We show that a trivial machine-learning model can achieve strong performance on existing benchmarks despite entirely discarding the information contained within MS/MS spectra themselves, and without using any other auxiliary information. This performance arises because compounds with reference MS/MS spectra are structurally distinct from those found in generic chemical databases, and machine-learning models can exploit this dissimilarity by learning to predict whether a compound is likely to have been measured by MS/MS. However, we show that this confound can be overcome by using a generative model to sample decoy structures that are chemically indistinguishable from compounds in reference MS/MS libraries. The resulting benchmark cannot be solved without learning from MS/MS spectra themselves. We leverage this benchmark to compare 17 published machine-learning models for MS/MS annotation, and find that many of these models fail to outperform simple baselines and may learn little about MS/MS itself. In contrast, a subset of models show convincing evidence of generalization. Our work provides a sound foundation for developing and evaluating computational methods for small molecule MS/MS.

---

High-throughput techniques can now reliably measure the DNA, RNA, and protein content of any biological sample, but comprehensively characterizing the small molecules within biological systems has proven more challenging. Liquid chromatography-mass spectrometry (LC-MS) routinely detects thousands of small molecule-associated signals in biological samples, but the majority cannot be matched to known compounds^1,2^. Structural annotation of these signals relies on tandem mass spectrometry (MS/MS), in which individual analytes are fragmented and the mass-to-charge ratios of the resulting fragments are recorded^3^. MS/MS spectra are commonly annotated *via* comparisons to libraries of reference MS/MS spectra from chemical standards^4–7^. However, typically only a fraction of MS/MS spectra can be annotated in this manner, in part because spectral libraries are small and are generally biased towards well-studied compounds for which chemical standards are commercially available^8,9^. Expert chemists can often deduce the presence and arrangement of key substructures based on manual interpretation of MS/MS spectra, but this is a time-consuming and labour-intensive process that does not scale to the thousands of spectra acquired in a typical metabolomic experiment^10^. Accordingly, the possibility of automating the interpretation of mass spectral data using computational approaches has been of longstanding interest, dating back to the DENDRAL project in the 1960s^11^. Interest in this problem has been reinvigorated over the past decade, culminating in the introduction of dozens of machine-learning models that aim to identify small molecules from their MS/MS spectra, even when these compounds do not appear in any reference spectral library^12–33^.

With the rapid proliferation of machine-learning models for MS/MS interpretation, the question of how to compare the performance of these models has received increasing attention^10,34–40^. However, meaningful comparison of computational methods for MS/MS interpretation has proven surprisingly difficult. New machine-learning models have historically been trained and evaluated on different datasets, using different evaluation set-ups and different metrics, and benchmarked selectively against relevant baselines—generally by authors who are proposing a new model and are therefore motivated to show that their model outperforms these baselines (the so-called “continental breakfast included effect”^41,42^). All of these factors can make it difficult to discern whether a particular model reliably outperforms other models and, if so, which specific technical decisions account for that improvement. This epistemological limitation has motivated the development of standardized benchmarks to enable more objective evaluation of model performance^34,37,39,40^. However, the validity of these benchmarks as measures of MS/MS interpretive ability has not been systematically examined. As a result, it remains unclear whether reported improvements in performance on these benchmarks translate to meaningful advances in practice.

Here, we expose a fundamental conceptual flaw in the evaluation of computational methods for MS/MS interpretation. We show that a trivial machine-learning model can achieve state-of-the-art performance on existing benchmarks despite wholly discarding the information contained within MS/MS spectra themselves, and without using any other auxiliary information. We elucidate the mechanism underlying this performance, finding that compounds with reference MS/MS spectra are structurally distinct from those found in generic chemical databases such as PubChem, and that machine-learning models can exploit this dissimilarity by learning to predict whether or not a compound is likely to have been measured by MS/MS rather than learning the chemical principles of small molecule fragmentation. The implication is that existing approaches for evaluating computational methods for MS/MS interpretation are spurious: that is, they do not measure aspects of model performance that they are widely assumed to measure. However, we show that the spurious nature of these evaluations can be overcome by leveraging a generative model to produce decoy structures that resemble those of compounds found within reference MS/MS libraries. The resulting benchmark cannot be solved without learning from MS/MS spectra, and therefore provides an epistemologically valid framework to evaluate the degree to which machine-learning models learn the complex mapping between spectra and structures^43^. Finally, we leverage this dataset to carry out the largest comparison of computational methods for MS/MS interpretation to date, encompassing 17 published models. Together, these results provide a sound foundation to systematically compare and optimize machine-learning approaches for structural annotation of small molecule MS/MS spectra.

## Results

### A fundamental confound in the evaluation of computational methods for small molecule MS/MS

Computational methods to interpret MS/MS spectra have generally been evaluated according to a standard conceptual framework that is designed to mimic their real-world deployment (**Fig. 1a**). In this framework, a method is presented with an MS/MS spectrum and a list of candidate structures, and is tasked with identifying the structure that generated the spectrum in question (hereafter, the ground-truth structure). Candidate structures are typically retrieved by filtering generic chemical structure databases such as Pub-Chem based on the exact mass or molecular formula of the ground-truth structure^9^. The number of candidate structures for each spectrum is typically large (on the order of hundreds to thousands), because databases such as PubChem contain far more unique structures than are present in reference MS/MS libraries. This surplus can be attributed at least in part to the fact that many compounds deposited in PubChem are unlikely to be present in biological systems, but instead represent synthetic compounds made in the laboratory or hypothetical structures explored as part of virtual screening campaigns^44^.

**Fig. 1.**
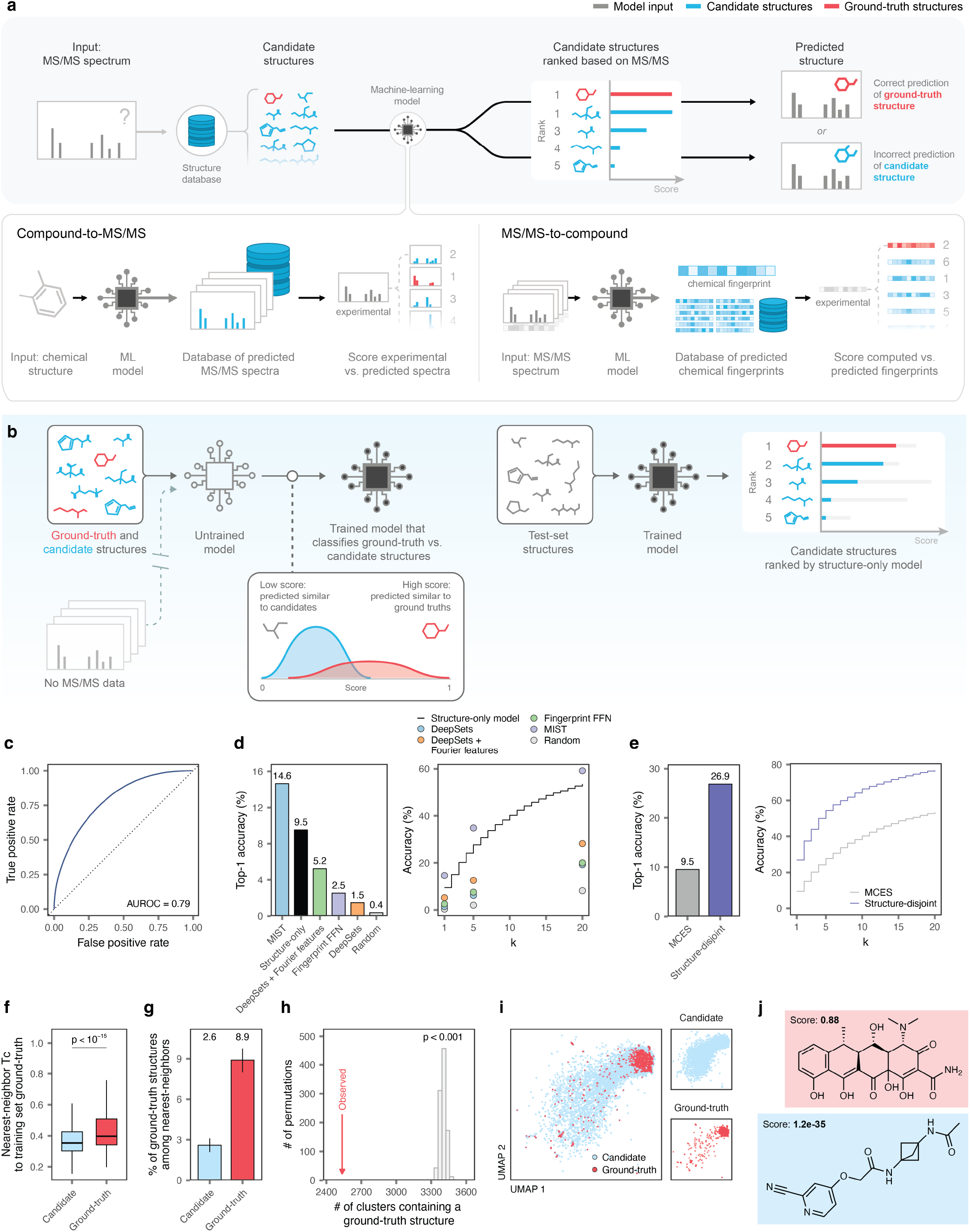
Spurious evaluations of computational methods for MS/MS interpretation. **a**, Schematic overview of existing approaches to benchmark computational methods for the annotation of MS/MS spectra on the basis of their ability to retrieve the correct structure from a set of candidate structures. **b**, Schematic overview of our “structure-only” model, which differentiates ground-truth from candidate structures while entirely discarding MS/MS spectra themselves. **c**, Receiver operating characteristic (ROC) curve showing the performance of the structure-only model in distinguishing between ground-truth and candidate structures in the held-out test fold of MassSpecGym. Inset text shows the area under the ROC curve (AUROC). **d**, Top-1 accuracy, left, or top-*k* accuracy (for *k* ≤ 20), right, of the structure-only model, as compared to baselines reported in MassSpecGym. **e**, Top-1 accuracy, left, or top-*k* accuracy (for *k* ≤ 20), right, of the structure-only model, when splitting spectra and candidate sets in MassSpecGym by MCES versus a more conventional structure-disjoint split. **f**, Nearest-neighbor Tanimoto coefficient to a ground-truth structure in the training set, for ground-truth versus candidate structures in the held-out test set of MassSpecGym. **g**, Proportion of ground-truth versus candidate structures in the held-out test set of MassSpecGym whose nearest-neighbors in the training set are themselves ground-truth structures. **h**, Number of clusters that contain at least one ground-truth structure, when clustering a large random sample of ground-truth and candidate structures from MassSpecGym, as compared to the number of clusters containing at least one ground-truth structure when cluster assignments are permuted. **i**, UMAP visualization of the chemical space occupied by ground-truth and candidate structures from 1,000 randomly sampled MassSpecGym candidate sets, with the former superimposed over the latter. **j**, Examples of test set compounds predicted to be ground-truth or candidate structures with high confidence by the structure-only model.

Whereas PubChem includes many structures that do not occur in nature, reference MS/MS libraries are typically biased towards compounds of biological interest. Cheminformatic analyses have repeatedly shown that certain physicochemical properties and substructures are enriched among biogenic compounds, as compared to synthetic compounds^45–47^. We reasoned that if the ground-truth compounds that have been measured by MS/MS could be distinguished from candidates drawn from PubChem based on these structural features, then these differences might confound the evaluation of computational methods for MS/MS interpretation. Specifically, we hypothesized that machine-learning models might learn to identify structures that are likely to have been measured by MS/MS, rather than learning the underlying chemical principles of small molecule fragmentation. In the extreme, we conjectured that a model could achieve competitive performance on existing benchmarks while disregarding MS/MS spectra altogether.

We tested this hypothesis in MassSpecGym, a recently introduced benchmark for MS/MS interpretation^34^. We assembled the structures that had been measured by MS/MS and their corresponding candidates from the training set of MassSpecGym, and trained a graph neural network to differentiate ground-truth from candidate structures; we refer to this classifier as the structure-only model (**Fig. 1b**). In the held-out test set, the structure-only model accurately classified structures as ground-truth or candidates, achieving an accuracy of 99.6% and an area under the receiver operating characteristic curve (AUROC) of 0.79 (**Fig. 1c**). This accuracy in binary classification translated into competitive retrieval of the ground-truth structure from among a large set of candidates: the structure-only model correctly retrieved the ground-truth structure in 9.5% of candidate sets, and ranked the ground-truth structure within its top-five predictions in 27.7% of candidate sets (top-20, 53.5%; **Fig. 1d** and **Supplementary Fig. 1a**). For comparison, the best model evaluated by the authors of MassSpecGym retrieved the correct structure in 14.6% of candidate sets. On the other hand, our structure-only model outperformed all four of the other models benchmarked by the authors of MassSpecGym. In the bonus challenge of MassSpecGym, in which candidate sets were defined based on the molecular formula of the ground-truth structure rather than its exact mass, our structure-only model achieved apparently state-of-the-art performance (**Supplementary Fig. 1b-c**). The performance of the structure-only model was slightly diminished when ground-truth and candidate structures were reprocessed to ensure identical standardization of tautomers, but its rank relative to published baselines was unchanged (**Supplementary Fig. 1d**). Thus, a trivial machine-learning model can achieve strong performance on existing benchmarks for MS/MS interpretation despite entirely ignoring the information provided within MS/MS spectra themselves.

The performance of certain computational methods in MassSpecGym is lower than had been reported in previous evaluations. This discrepancy likely reflects the stringent split of the training and test folds in MassSpecGym, whereby the held-out test set contains structures with a minimum bond edit distance of at least 10 to any compound in the training set^48^. Historically, the more common approach has been that of structure-disjoint cross-validation, in which all MS/MS spectra from a given compound are assigned to the same fold, while permitting chemically related structures to be partitioned into different folds. With a more conventional structure-disjoint split, the performance of the structure-only model was even higher, with the ground-truth structure now correctly retrieved in 26.9% of candidate sets (top-5, 54.5% and top-20, 76.9%; **Fig. 1e** and **Supplementary Fig. 1e**). Thus, under the structure-disjoint splits that were conventional prior to MassSpecGym, a trivial model can appear to be remarkably useful for small molecule identification.

We sought to clarify the mechanism that allowed our structure-only model to achieve excellent performance in the annotation of MS/MS spectra, despite disregarding the information contained within these spectra altogether. We hypothesized that ground-truth structures in the training and test folds shared structural features that differentiated them from candidate structures from PubChem, despite the stringent bond edit distance threshold used to define this split. A series of analyses corroborated this hypothesis. First, we quantified the chemical similarity between the structures in the held-out test fold of MassSpecGym and the structures in the training folds, to which the structure-only model had been exposed. Ground-truth structures in the test fold exhibited significantly greater chemical similarity to ground-truth structures in the training folds than to candidate structures (**Fig. 1f**). Accordingly, the nearest-neighbors of ground-truth structures in the test fold were significantly enriched for ground-truth structures in the training folds, relative to candidate structures (**Fig. 1g**). Moreover, when clustering a random subset of ground-truth and candidate structures in MassSpecGym, we observed that ground-truth structures were found in significantly fewer clusters than random expectation (**Fig. 1h**). Finally, we visualized the chemical space occupied by ground-truth and candidate structures using the nonlinear dimensionality reduction algorithm UMAP, and found that ground-truth structures were sparsely distributed within this chemical space (**Fig. 1i**). We show examples of unseen compounds predicted to be ground-truth or candidate structures with high confidence by the structure-only model in **Fig. 1j** and **Supplementary Fig. 1f**.

Together, these results demonstrate that it is possible to achieve competitive performance on existing benchmarks for the interpretation of small molecule MS/MS despite wholly discarding the information contained within MS/MS spectra themselves. This performance reflects the ability of machine-learning models to distinguish the small molecules for which reference MS/MS have been acquired from those contained in generic chemical structure databases such as PubChem. Importantly, this phenomenon is distinct from the problem of ‘meta-scores,’ whereby incorporation of auxiliary data such as citation counts or production volumes into machine-learning models biases these models towards well-studied compounds^49,50^, and from the related observation that ranking candidates by the order in which they are returned by the PubChem API often recovers the ground-truth structure^51^. Here, we show that a trivial machine-learning model can make accurate predictions by learning solely from chemical structures, without using any auxiliary information whatsoever.

### Spurious performance persists in an expanded benchmark dataset

The competitive performance of the structure-only model in existing benchmarks for the interpretation of small molecule MS/MS spectra suggests that the prevailing approach to model evaluation does not quantify aspects of model performance that it purports to. We therefore set out to develop a benchmark that would evaluate the ability of machine-learning models to interpret MS/MS spectra, rather than rewarding the recognition of chemical structures likely to appear in reference spectral libraries.

We initially speculated that the performance of our structure-only model might have arisen in part from the design of MassSpecGym. In MassSpecGym, candidate sets for each spectrum are limited to a maximum of 256 structures, which is considerably fewer than would typically be considered when searching generic structure databases such as PubChem. Moreover, we found that the effective number of structures in any given candidate set was sometimes substantially less than 256—for instance, because the same chemical structures appeared in the same candidate set multiple times as distinct tautomers, stereoisomers, or isotopomers, or because the candidate set contained implausible structures, such as those composed of multiple disconnected fragments or with extreme formal charges (**Supplementary Fig. 2a-m**). Removal of duplicate and implausible structures further decreased the total number of candidates, and therefore the difficulty of the benchmarking task (**Supplementary Fig. 2n-o**). We hypothesized that the structure-only model would struggle to identify ground-truth structures within greatly expanded candidate sets.

To test this hypothesis, we turned from MassSpecGym to Spectraverse, a more comprehensive dataset of reference MS/MS spectra that we recently introduced^36^. We retrieved candidate sets from PubChem for each spectrum based on either the exact mass or the molecular formula of the ground-truth structure (**Fig. 2a**). The resulting candidate sets were, on average, 22.4 and 6.8 times larger than those in MassSpecGym for mass and formula candidates, respectively (**Fig. 2b** and **Supplementary Fig. 3a-d**).

**Fig. 2.**
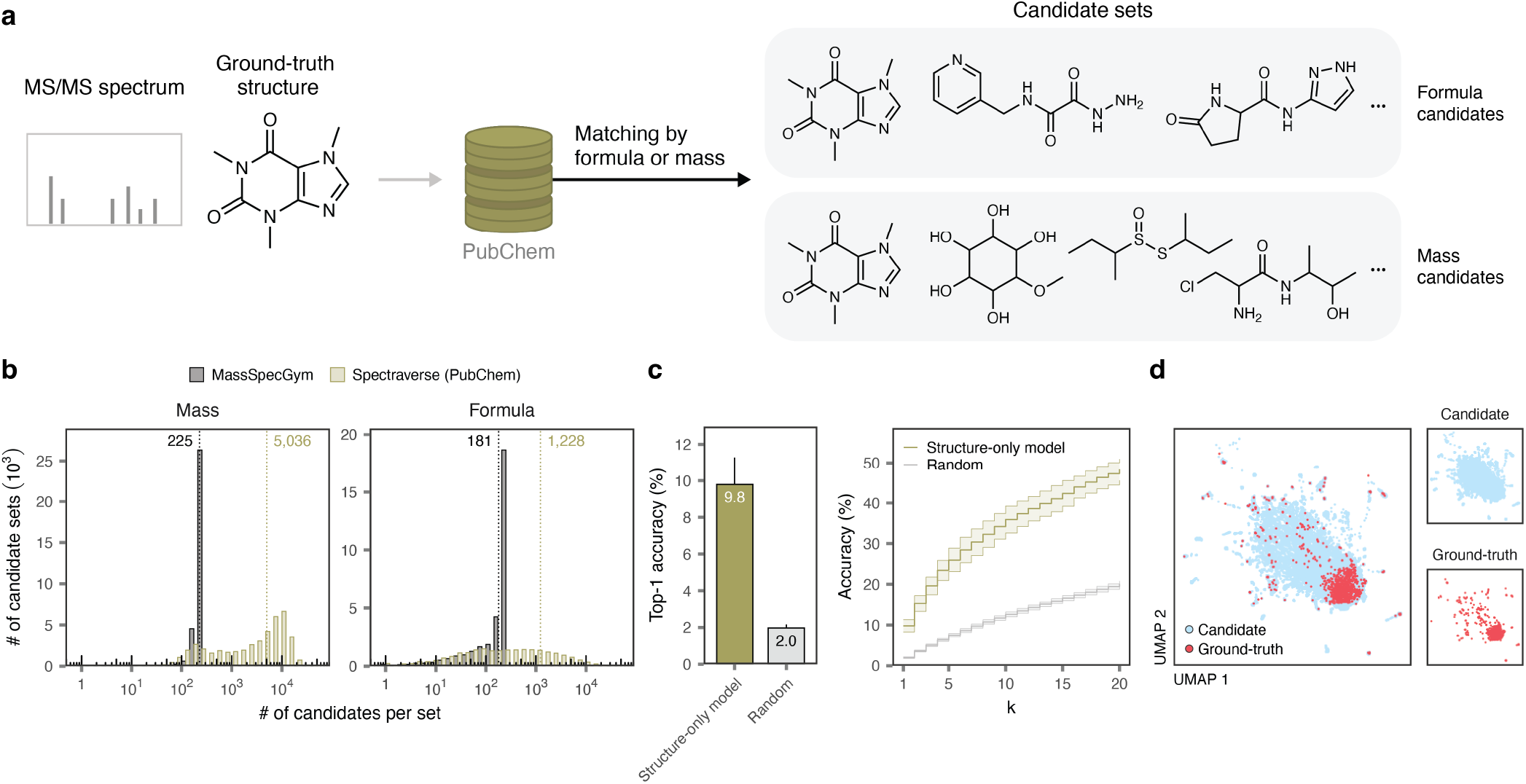
Spurious performance in a markedly expanded benchmark dataset. **a**, Schematic overview of PubChem candidate set creation for MS/MS spectra in Spectraverse. **b**, Number of candidate structures associated with each MS/MS spectrum in MassSpecGym versus expanded candidate sets retrieved from PubChem for MS/MS spectra in Spectraverse, shown separately for mass-versus formula-based candidate sets. **c**, Top-1 accuracy, left, or top-*k* accuracy (for *k* ≤ 20), right, of the structure-only model in expanded PubChem candidate sets, as compared to a random selection baseline, for formula-based candidate sets in structure-disjoint cross-validation. Error bars show standard deviation across ten cross-validation folds. **d**, UMAP visualization of the chemical space occupied by ground-truth and candidate structures from *n* = 1,000 randomly selected PubChem candidate sets, with the former superimposed over the latter.

We then evaluated the performance of the structure-only model in these expanded candidate sets. Our expectation was that, with the average size of each candidate set increased by an order of magnitude or more, it would no longer be possible to identify the ground-truth structure without access to MS/MS data. To our surprise, however, the structure-only model continued to accurately retrieve the ground-truth structure from among a large set of candidates, now identifying the correct structure in 9.8% of formula-based candidate sets and 6.1% of mass-based candidate sets (**Fig. 2c** and **Supplementary Fig. 3e**). The performance of the structure-only model decreased when training and test folds were separated by a minimum MCES distance threshold, but continued to substantially outperform a random baseline (**Supplementary Fig. 3f-g**). As in MassSpecGym, UMAP visualization revealed that the ground-truth structures in Spectraverse were concentrated in chemically distinct subregions within the broader chemical space of candidate structures (**Fig. 2d**).

Thus, compounds that have been measured by MS/MS can be differentiated from thousands of isomeric or isobaric candidates on the basis of their structures alone. As a result, greatly increasing the number of candidate structures associated with each spectrum fails to address a more fundamental confound in the longstanding paradigm by which methods for the interpretation of small molecule MS/MS spectra have been evaluated.

### Generating candidate structures with a chemical language model

The competitive performance of the structure-only model reflects its ability to differentiate compounds that have been measured by MS/MS from candidate structures drawn from PubChem, which are structurally distinct. We reasoned that if candidate sets were instead populated with structures drawn from the same regions of chemical space as the compounds in reference MS/MS libraries, then this trivial separation would no longer be possible. This would be desirable because the relative performance of machine-learning models in the resulting benchmark would then more directly reflect the ability of these models to learn the complex mapping between spectra and structures. We therefore set out to construct such a benchmark.

We hypothesized that a generative model trained on compounds in reference MS/MS libraries could produce candidate structures that could not be trivially differentiated from ground-truth structures. Chemical language models have emerged as a particularly powerful class of generative models for exploring the chemical space around a training set of known molecules^52–54^. These models represent chemical structures as short strings of text, and leverage neural network architectures originally developed for natural language processing to learn the statistical properties of these strings. Prior work has established that chemical language models can generate novel structures that closely resemble the structures in their training sets^55–58^. We therefore trained a chemical language model on all ground-truth structures with reference MS/MS spectra in Spectraverse, and used this model to generate a large library of novel structures (**Fig. 3a**). We then constructed mass- and formula-based candidate sets for each MS/MS spectrum based on this generated library (**Supplementary Fig. 4a-f**).

**Fig. 3.**
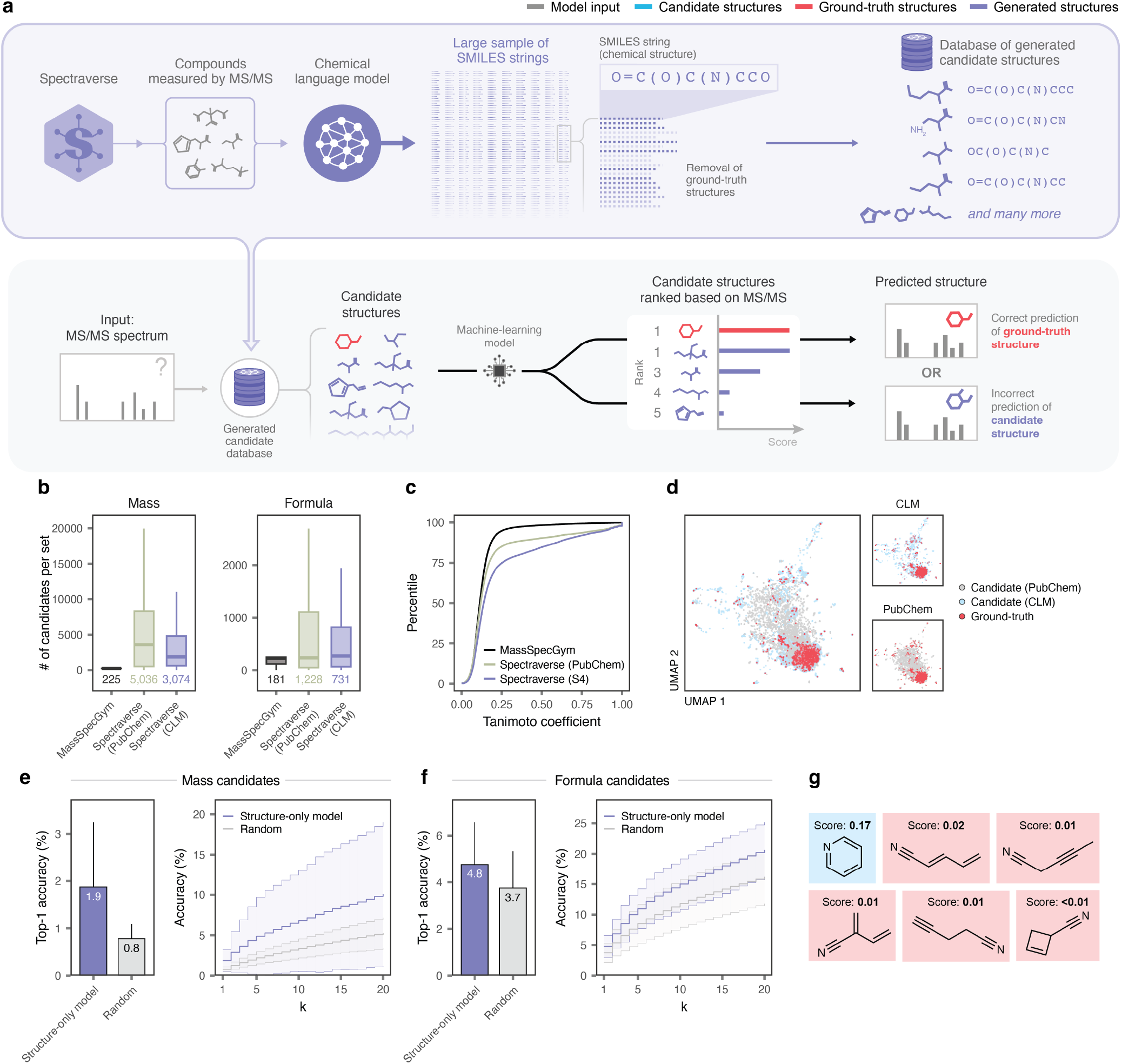
Generating candidate structures with a chemical language model. **a**, Schematic overview of candidate structure generation with a chemical language model trained on structures with reference MS/MS spectra in Spectraverse. **b**, Number of PubChem-derived or language model-generated candidate structures associated with each MS/MS spectrum in Spectraverse, shown separately for mass-versus formula-based candidate sets. **c**, Cumulative distribution function of Tanimoto coefficients between ground-truth structures and one randomly selected structure per candidate set, for MassSpecGym, PubChem-derived candidate sets, or language model-generated mass-based candidate sets. **d**, UMAP visualization of the chemical space occupied by ground-truth and PubChem-derived or language model-generated candidate structures from *n* = 1,000 randomly selected candidate sets, with the former superimposed over the latter. **e**, Top-1 accuracy, left, or top-*k* accuracy (for *k* ≤ 20), right, of the structure-only model in language model-generated candidate sets, as compared to a random selection baseline, for mass-based candidate sets with a MCES distance split. Error bars and shaded areas show standard deviation across ten cross-validation folds. **f**, As in **e**, but for formula-based candidate sets. **g**, Example of a generated candidate set in which the ground-truth compound can be deduced by the structure-only model without access to MS/MS information.

Generated candidate sets were comparable in size to PubChem-derived candidate sets, and considerably larger than those in MassSpecGym (**Fig. 3b**). However, language model-generated candidates were more structurally similar to ground-truth structures than candidates from either PubChem or MassSpecGym (**Fig. 3c-d** and **Supplementary Fig. 4g-i**). As a result, the structure-only model failed to reliably differentiate ground-truth structures from generated candidates without access to MS/MS information. In mass-based candidate sets, its top-1 accuracy of 1.9% was only marginally higher than that of a random baseline, at 0.8% (**Fig. 3e**). Performance was higher in formula-based candidate sets, but remained only marginally above chance (**Fig. 3f**).

Although the performance of the structure-only model was greatly diminished in generated candidate sets, it remained slightly higher than random expectation. To understand why this was the case, we inspected the candidate sets for which the structure-only model made accurate predictions. We found that, for many of these candidate sets, only a single candidate appeared structurally plausible, with all other generated structures representing chemically implausible or unstable isomers (**Fig. 3g** and **Supplementary Fig. 4j**). Thus, for a small subset of MS/MS spectra, generative models cannot generate realistic decoy structures, and the ground-truth structure can be deduced without requiring MS/MS data.

Together, these results demonstrate that generative models can sample candidate structures from the same regions of chemical space as compounds in reference MS/MS libraries. The resulting benchmark cannot be solved without learning from MS/MS spectra, and therefore provides a sound foundation to evaluate computational methods for MS/MS interpretation.

### Benchmarking computational methods for MS/MS interpretation

With these candidate sets in hand, we sought to benchmark the performance of computational methods for structural annotation of MS/MS spectra. Although computational methods for MS/MS interpretation have previously been evaluated in a variety of settings, we recognized that Spectraverse offered an opportunity for a more rigorous comparison than had previously been possible because it addresses a number of limitations of existing benchmarks, such as MassSpecGym or the CASMI challenges^34,39,59^. First, and most importantly, ground-truth structures cannot be trivially differentiated from candidate structures without learning from MS/MS information, implying that any observed differences in performance reflect true differences in the capacity of machine-learning models to interpret MS/MS spectra. Second, Spectraverse provides a large and diverse dataset comprising >480,000 spectra from >44,000 compounds for model training and evaluation, which should increase the degree to which estimates of model performance generalize to unseen spectra. Third, Spectraverse contains MS/MS spectra from a diverse range of ionization modes and adducts that are commonly encountered in untargeted metabolomics experiments. Fourth, Spectraverse specifies cross-validation folds rather than a fixed train/test split, allowing for statistically rigorous comparisons of model performance rather than relying on comparisons of point estimates. Fifth, Spectraverse explicitly specifies the data to be used for model training in order to isolate the effects of modelling decisions from the composition of the training dataset, which we regard as particularly important given the well-documented pitfalls of incorporating auxiliary datasets in machine-learning models for MS/MS interpretation^49,50^. Sixth, Spectraverse explicitly specifies candidate structures for each spectrum rather than leaving candidate retrieval to individual researchers, thereby ensuring that all models are evaluated under identical conditions. Finally, the relatively large size of the language model-generated candidate sets is commensurate with the number of candidates typically retrieved from generic chemical databases such as PubChem. This provides a practical test of computational efficiency, in that models that are unable to make predictions at this scale are unlikely to find practical use in searching those databases.

We leveraged our benchmark to compare 17 published methods for structural annotation of MS/MS spectra from small molecules. Historically, machine-learning models have taken one of two broad approaches to learn the mapping between MS/MS spectra and chemical structures. One class of models takes a chemical structure as input and predicts the corresponding MS/MS spectrum. Another major class of models takes a MS/MS spectrum as input and predicts certain physicochemical or structural properties of the corresponding small molecule, often in the form of a chemical fingerprint. More recently, new approaches have been introduced that seek to co-embed chemical structures and MS/MS spectra in a shared latent space using approaches such as contrastive learning. We benchmarked a range of models from each of these categories, focusing on approaches that provided open-source code for model training and inference. We also included several baselines, including random selection, the structure-only model, and a recently proposed baseline in which each test spectrum is assigned the chemical fingerprint of the ground-truth structure associated with its nearestneighbor spectrum in the training set^60^. We implemented this nearest-neighbor baseline using three measures of spectral similarity, including cosine, neutral loss cosine, and modified cosine similarity^61^.

Surprisingly, most of the 17 published models that we benchmarked failed to outperform one or more of these baselines (**Fig. 4a-d** and **Supplementary Fig. 5a-d**). Seven of these models did not even significantly outperform the structure-only baseline, suggesting that the previously reported performance of these models reflects their ability to differentiate ground-truth from candidate structures rather than their ability to interpret MS/MS spectra *per se*. Moreover, 13 of the 17 models failed to outperform the modified cosine variant of the nearest-neighbor baseline, and only FraGNNet and ICEBERG achieved a significantly higher top-1 accuracy. FraGNNet was the top-performing method overall, achieving a top-1 accuracy of 20.1% in formula-based candidate sets (17.2% in mass-based candidate sets), whereas ICEBERG achieved top-1 accuracies of 14.1% and 9.4%, respectively. Although the top-1 accuracy of these models was modest, both were often able to rank the ground-truth structure within a short list of candidates, achieving top-10 accuracies of 43.0% and 36.2% of formula-based candidate sets, respectively (**Fig. 4b** and **Supplementary Fig. 5b,e**).

**Fig. 4.**
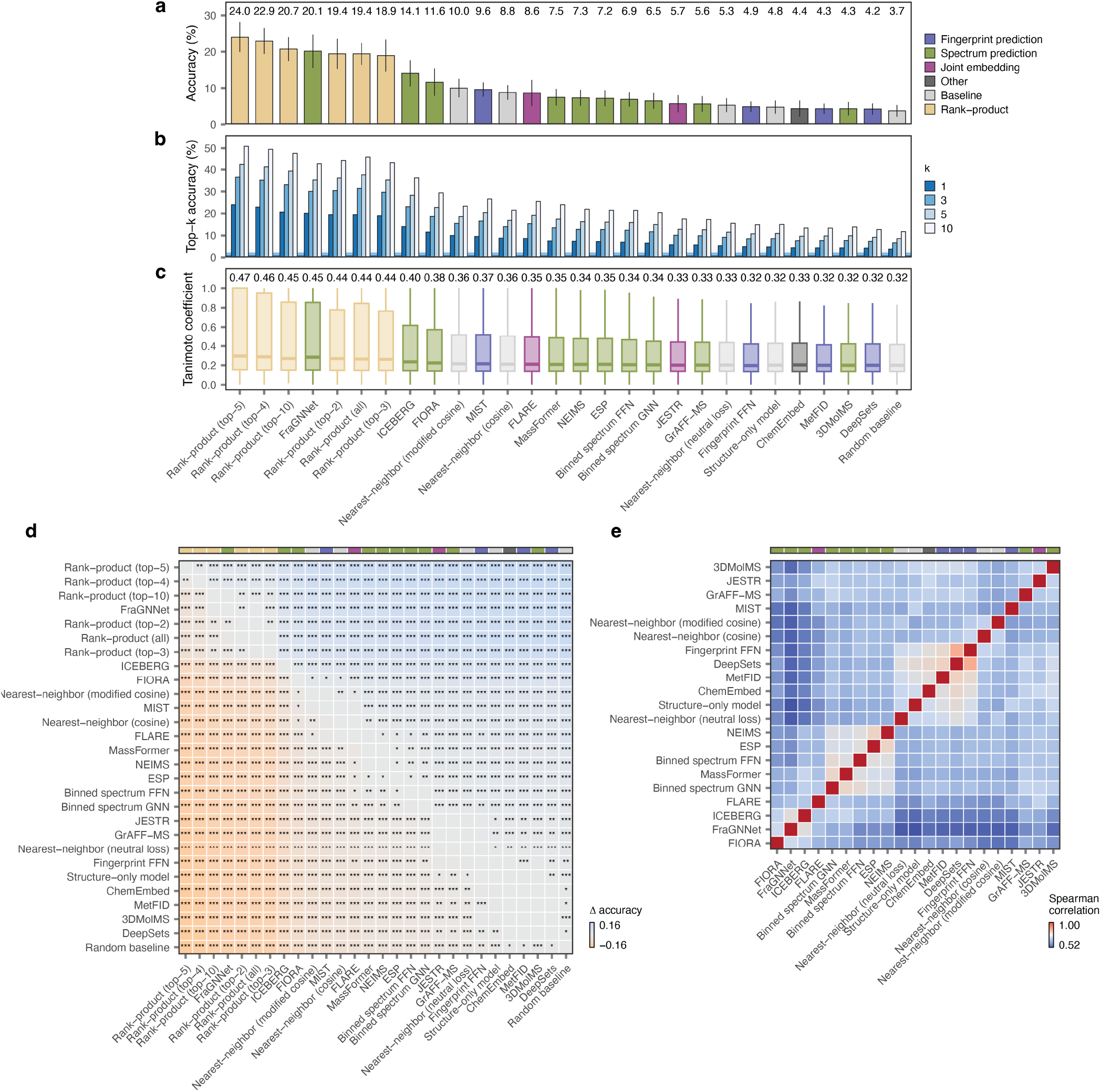
An unconfounded benchmark of computational methods for MS/MS annotation. **a**, Top-1 accuracy of 17 published models for MS/MS annotation, baseline approaches, and rank-product aggregation of varying numbers of top-performing models, in formula-based candidate sets. Error bars show standard deviation across ten cross-validation folds. Inset text shows the mean top-1 accuracy across folds. **b**, As in **a**, but showing the top-3, top-5, and top-10 accuracy for each model. **c**, As in **a**, but showing the Tanimoto coefficient between the ground-truth structure and the top-ranked candidate. Inset text shows the mean Tanimoto coefficient. **d**, Mean difference in the top-1 accuracy (Δ accuracy) between the models shown in **a**. Text highlights comparisons with a two-sided paired t-test p-value less than 0.05 (*, p < 0.05; **, p < 0.01; ***, p < 0.001). **e**, Spearman correlation matrix between ranks assigned to ground-truth structures by the 17 published models for MS/MS annotation shown in **a**.

We also asked whether, beyond identifying individual top-performing models, our benchmark identified more general principles for accurate annotation of MS/MS spectra. Notably, because every model was evaluated on the same set of MS/MS spectra, our data permitted a direct comparison of spectrum prediction and fingerprint prediction approaches. While the two best-performing models sought to predict MS/MS spectra from chemical structures, many other spectrum prediction models performed poorly, often under-performing the modified cosine nearest-neighbor baseline or even the structure-only model. The same pattern held for fingerprint prediction and joint embedding approaches, some of which performed only marginally better than random, while others were among the top-performing models. We also found no significant correlation between performance and model size, as measured by the number of parameters (**Supplementary Fig. 5f**). Thus, our data do not clearly favor any one paradigm for MS/MS annotation but instead suggest that more specific aspects of model design and training ultimately determine model performance.

We found that the ranks assigned to ground-truth structures by each of the 17 models we benchmarked were weakly correlated, suggesting that these models capture complementary aspects of the mapping between MS/MS spectra and structures (**Fig. 4e** and **Supplementary Fig. 5g**). In other areas of computational biology, past benchmarks have found that, when many methods are available for a given task, aggregating predictions across models often yields more accurate results than any individual method (the so-called “wisdom of the crowds” effect)^62,63^. To test whether the same principle extends to small molecule MS/MS, we used a simple rank-product aggregation strategy to combine predictions made by various subsets of top-performing models. We found that aggregating the top-5 models (FraGNNet, ICEBERG, FIORA, MIST, and FLARE) afforded the highest performance and improved substantially over FraGNNet alone, achieving a top-1 accuracy of 24.0% and a top-10 accuracy of 50.8% in formula-based candidate sets (**Fig. 4a-d** and **Supplementary Fig. 5a-d**).

Together, these results provide a large-scale comparison of computational methods for MS/MS annotation in a setting in which ground-truth and candidate structures cannot be trivially differentiated without attending to the MS/MS spectra themselves. We find that (1) most published models do not outperform simple baselines, and many do not even outperform a trivial structure-only model, suggesting that their performance on past benchmarks can be wholly attributed to their ability to recognize structures likely to have been measured by MS/MS; (2) two models, FraGNNet and ICEBERG, significantly outperform these baselines and show evidence of generalizing beyond their training sets; and (3) existing models are complementary, such that combining predictions from multiple models can materially improve performance.

## Discussion

In this work, we expose a pernicious conceptual flaw in the evaluation of computational methods for structural annotation of MS/MS spectra from small molecules. We show that compounds in reference MS/MS libraries are sufficiently distinct from those that populate generic chemical databases such as PubChem that a trivial model—one that wholly disregards MS/MS spectra themselves—can achieve strong performance on existing benchmarks for MS/MS interpretation, in some cases outperforming well-regarded models. Our trivial model correctly identifies the ground-truth structure for 9.5% of the MS/MS spectra in MassSpecGym, and its top-1 accuracy rises to 13.8% when the molecular formula is provided and to 26.9% under a more conventional structure-disjoint split. Importantly, this phenomenon is not an artifact of MassSpecGym, since this structure-only model continues to demonstrate strong performance in PubChem-derived candidate sets that are on average an order of magnitude larger. This finding implies that existing approaches to model evaluation in small molecule MS/MS are spurious: that is, a model’s performance need not reflect its ability to learn the underlying chemical principles of small molecule fragmentation. However, we demonstrate that this confound can be eliminated by using a chemical language model to sample candidate structures from the same regions of chemical space as the ground-truth structures themselves. The resulting benchmark abolishes the spurious performance of a structure-only model and yields an epistemologically valid framework to evaluate the capacity of machine-learning models to learn the mapping between spectra and structures. We leverage this dataset to carry out the largest benchmark of computational methods for MS/MS interpretation to date, comparing the performance of 17 published models encompassing spectrum prediction, fingerprint prediction, and joint embedding approaches.

The possibility of automating the identification of small molecules from their MS/MS spectra has been of longstanding interest. A renaissance of interest in this problem over the past decade has led to the development of a series of increasingly complex machine-learning models for MS/MS interpretation. Why has this proliferation of machine-learning methods yet to translate into a marked increase in metabolite annotation rates in experimental datasets? One possibility that has long been recognized is that the difficulty in achieving objective comparisons of these models has hindered progress, insofar as it has not been straightforward to elucidate the impact of specific modelling decisions on the accuracy of these models^10^. Our work highlights a deeper issue: even apparently objective comparisons of these models may not actually measure aspects of model performance that they are widely assumed to. In fact, existing benchmarks are so confounded that a model without any access to MS/MS spectra can achieve performance that is competitive with well-regarded models. Our findings might explain, at least in part, why reliable annotation of small molecule MS/MS spectra remains challenging: efforts to optimize model performance on existing benchmarks may have misguided developers by rewarding models for recognizing structures likely to have been measured by MS/MS, rather than the ability to interpret MS/MS spectra *per se*.

The availability of this epistemologically valid benchmark afforded an opportunity to evaluate the performance of 17 published models for structural annotation of MS/MS spectra in an unconfounded setting. The most striking finding to emerge from this comparison is that, when these models cannot exploit differences in the properties of ground-truth versus candidate structures, their performance is markedly lower than has been previously reported. Seven of the 17 models failed to significantly outperform the structure-only model, which is consistent with the possibility that these models rely on the distinctive features of ground-truth structures for MS/MS annotation and learn comparatively little about MS/MS spectra themselves. Moreover, only two models, FraGNNet and ICEBERG, significantly outperformed a more stringent baseline in which each test spectrum was assigned the chemical fingerprint associated with its nearest-neighbor in the training set (as quantified by the modified cosine similarity). While the top-1 accuracy of both FraGNNet and ICEBERG remained modest, both models were often able to rank the correct structure within a short list of candidates, even in this highly challenging setting. These and other top-performing models also proved complementary to one another, such that performance substantially higher than FraGNNet alone could be achieved through a simple rank-product aggregation of the top-5 models (FraGNNet, ICEBERG, FIORA, MIST, and FLARE). Interestingly, both top-performing models adopt broadly similar strategies aiming to explicitly model the process by which small molecules fragment within the mass spectrometer, suggesting that this may be a particularly promising direction for future work. The strong performance of our rank-product aggregation approach, however, also suggests that future efforts should seek to leverage the apparently complementary strengths of all three paradigms studied here (i.e., fingerprint prediction, spectrum prediction, and joint embedding) to maximize performance.

We have made Spectraverse, including all MS/MS spectra, generated candidate sets, and crossvalidation fold assignments, available from Zenodo at https://doi.org/10.5281/zenodo.19927403, and offer several recommendations for its use. First, models should be trained exclusively on the ground-truth structures and MS/MS spectra provided in Spectraverse itself, so that differences in model performance can be attributed to modelling decisions rather than the composition of the training dataset or the use of auxiliary data. Second, results should be reported across all ten cross-validation folds in order to permit statistical comparisons of model performance. Last, we encourage model developers to deposit the complete outputs of their models, including scores for every structure in each candidate set, in a public repository such as Zenodo so that the community may reproduce and independently scrutinize reported results.

## Methods

### Structure-only model

MassSpecGym candidate sets were obtained from HuggingFace (https://huggingface.co/datasets/roman-bushuiev/MassSpecGym, files MassSpecGym_retrieval_candidates_formula.json and MassSpec-Gym_retrieval_candidates_mass.json, both last updated August 20, 2024). Each candidate set consists of the ground-truth structure and a set of decoy molecules retrieved from a series of chemical structure databases by matching either the molecular formula or the mass of the ground-truth structure (i.e., formula and mass candidates, respectively).

Our structure-only model consisted of a message-passing neural network (MPNN), as implemented in the ChemProp package^64^. In this approach, chemical structures are represented as graphs, where atoms correspond to nodes and bonds to edges. ChemProp first constructs initial features based on the identity and properties of each atom or bond (for instance, atomic number, formal charge, or bond order). These initial features are then passed to an MPNN, and embeddings are updated over a series of message-passing steps. During message passing, each atom aggregates information from its neighboring atoms and bonds. Atom-level embeddings are then aggregated to produce a molecule-level embedding, which is fed through a feed-forward neural network to predict the molecular property of interest. Here, a ChemProp model was trained to perform binary classification of ground-truth versus candidate structures, using the MassSpecGym candidate sets exactly as distributed and with the same training/validation/test split. The model was trained for a maximum of 200 epochs, with early stopping triggered if the validation loss had not improved for 10 epochs.

To compute the top-*k* accuracy, candidate structures were ranked by the predicted ground-truth class probabilities from the structure-only model, and the proportion of candidate sets for which the ground-truth structure ranked among the top *k* candidates was calculated. The top-*k* accuracies shown in the figures for baseline methods were taken from the MassSpecGym paper^34^. These analyses were also repeated for a more conventional structure-disjoint split of MassSpecGym, in which ground-truth structures (and their corresponding candidate sets) were split into ten folds based on the first fourteen-character block of their InChIKeys. ROC curves and top-*k* accuracies shown in the manuscript represent the mean performance of ten independent training runs with different random seeds to account for stochasticity in model initialization and optimization.

We performed a series of additional analyses to rationalize the observed performance of the structure-only model. Each of the analyses described below was performed with the mass candidates. First, we evaluated the chemical similarity between ground-truth structures in the training and test sets of MassSpecGym. To this end, we randomly sampled 1,000 ground-truth structures and 1,000 candidate structures from the test fold of MassSpecGym, and an equal number of ground-truth and candidate structures from the training and validation folds combined, excluding any structures that also appeared in the test fold. Ground-truth and candidate structures were sampled in equal numbers in order to mitigate the propensity for comparisons of nearest-neighbor similarity to be confounded by the size of the chemical databases being compared^65,66^. For each of these structures, we computed a 2048-bit Morgan fingerprint using RDKit. Then, for each structure in the test set, we identified its nearest-neighbor among all of the structures in the training and validation folds, and recorded the corresponding Tanimoto coefficient (Tc). We compared the nearest-neighbor Tc for ground-truth versus candidate structures in the test fold. In addition, we computed the proportion of nearest-neighbors in the training and validation folds that were themselves ground-truth structures, separately for ground-truth versus candidate structures in the test fold. This latter analysis was performed with an increased number of candidate structures from the training and validation folds (1,000 ground-truth structures and 99,000 candidate structures) to better account for the class imbalance of the dataset.

To compare the coverage of chemical space by ground-truth and candidate structures within MassSpecGym, we next performed a clustering analysis. We randomly selected a total of 10,000 candidate sets (8,000, 1,000, and 1,000 from the training, validation, and test folds, respectively) and computed Morgan fingerprints for each unique structure, as described above. Then, these fingerprints were clustered using *k*-means with the MiniBatchK-Means function from scikit-learn (with *k* = 10,000; results were robust to the choice of *k*), and the number of clusters containing at least one ground-truth structure was computed. This value was then compared to the null expectation derived from 10,000 random permutations of cluster assignments.

Finally, all ground-truth and candidate structures in the test fold of MassSpecGym were embedded in two dimensions using UMAP, using the implementation in the umap-learn Python package with the number of neighbors (n_neighbors) set to 5,000 and the minimum distance (min_dist) set to 0.1.

### Analysis of MassSpecGym candidates

After obtaining MassSpec-Gym candidate sets from HuggingFace as described above, stereoisomers were identified by using the RDKit function MolToSmiles with arguments isomericSmiles=False, canonical=True to identify structures with the same canonical SMILES after removal of stereochemical information. Tautomers were identified by grouping structures with the same InChIKey. Fragments were identified as structures for which the RDKit function FragmentParent returned a structure with a mass at least 1.5 Da less than the original structure (to exclude mass differences that resulted solely from protonation or deprotonation of the original structure). Isotopologues were identified by iterating through the atoms in each structure and using the RDKit method SetIsotope to remove isotopic information. For the subset of structures whose canonical SMILES changed as a result, structures sharing the same first fourteen-character InChIKey block were labelled as isotopologues. Net charge was calculated using the RDKit function GetFormalCharge.

To count the total number of candidates remaining after removal of duplicate or implausible structures, candidate structures in each set were standardized according to the following pipeline. First, stereochemistry was removed with the function RemoveStereochemistry, after which the molecule was sanitized using the function SanitizeMol. Hydrogens were removed by first using the RemoveHs function and then reinstantiating the molecule object from its canonical SMILES, which we found to be necessary to remove some explicit hydrogens. Metal atoms were disconnected and the molecule reionized using the Cleanup function, and the largest fragment was identified and retained with the FragmentParent function, after which the molecule was again sanitized. Next, the molecule was converted to its canonical tautomer using the TautomerEnumerator class, and then sanitized again. The same pipeline was applied to the ground-truth structures to identify cases in which the ground-truth structure was present more than once in the candidate set. Duplicate structures (i.e., those with the same InChIKey) were removed, as were candidate structures with a mass that differed by at least 1.5 Da from that of the ground-truth structure.

### Spectraverse candidate sets

Because we found that candidate sets in MassSpecGym often contained redundant or implausible candidates, and because we found that simply querying PubChem with the chemical formula or monoisotopic mass of a compound with a reference MS/MS spectrum in Spectraverse sometimes failed to retrieve the ground-truth structure, we devised a multi-stage pipeline to generate expanded candidate sets for each MS/MS spectrum in Spectraverse. We began by downloading a total of 118.6 million chemical structures from Pub-Chem (file https://ftp.ncbi.nlm.nih.gov/pubchem/Compound/Extras/CID-SMILES.gz, downloaded September 8, 2024). We then standardized and deduplicated these structures, following an approach that we recently described^36^. In brief, SMILES strings were loaded in RDKit, sanitized, and cleaned to remove hydrogens, metals, and disconnected fragments using the RDKit functions SanitizeMol, Cleanup, RemoveHs, and FragmentPar-ent, respectively. Stereochemical information was removed, and tautomers were standardized to their canonical forms using the TautomerEnumerator class. Charges were neutralized when possible using a combination of the RDKit Uncharger class and the series of SMARTS patterns provided in the RDKit Cookbook. We found that this approach sometimes incorrectly neu-tralized functional groups that were written in charge-separated forms, no-tably sulfoxides and phosphoryl groups, and therefore implemented addi-tional checks to ensure that these functional groups were correctly neutral-ized. When neutralization altered the net charge of the molecule, both the charged and uncharged forms of the structure were retained for further anal-ysis. Structures containing radical electrons or atoms with invalid valences were removed. The output of this step was a set of deduplicated and stan-dardized two-dimensional structures from PubChem.

We next developed a pipeline to retrieve candidate structures based on the molecular formula of the ground-truth structure. We began by retrieving compounds whose molecular formulas exactly matched that of the ground-truth compound, but found that this approach sometimes failed to retrieve the ground-truth structure or other plausible isomers; moreover, when the same structure was represented in Spectraverse in two different charge states (for instance, as the [M+H]^+^ adduct of a zwitterion versus the [M]^+^ adduct of a compound with a net charge of +1), two different candidate sets would ject from its canonical SMILES, which we found to be necessary to remove sormepeoerxtpelidcirtehsyudlrtosg.ens. Metal atoms were disconnected and the molecule reionized using the Cleanup function, and the largest fragment was identified and retained with the FragmentParent function, after which the molecule was again sanitized. Next, the molecule was converted to its canonical tautomer using the TautomerEnumerator class, and then sanitized again. The same pipeline was applied to the ground-truth structures to identify cases in which the ground-truth structure was present more than once in the candidate set. Duplicate structures (i.e., those with the same InChIKey) were removed, as were candidate structures with a mass that differed by at least 1.5 Da from that of the ground-truth structure.

### Spectraverse candidate sets

Because we found that candidate sets in MassSpecGym often contained redundant or implausible candidates, and because we found that simply querying PubChem with the chemical formula or monoisotopic mass of a compound with a reference MS/MS spectrum in Spectraverse sometimes failed to retrieve the ground-truth structure, we devised a multi-stage pipeline to generate expanded candidate sets for each MS/MS spectrum in Spectraverse. We began by downloading a total of 118.6 million chemical structures from PubChem (file https://ftp.ncbi.nlm.nih.gov/pubchem/Compound/Extras/CID-SMILES.gz, downloaded September 8, 2024). We then standardized and deduplicated these structures, following an approach that we recently described^36^. In brief, SMILES strings were loaded in RDKit, sanitized, and cleaned to remove hydrogens, metals, and disconnected fragments using the RDKit functions SanitizeMol, Cleanup, RemoveHs, and FragmentParent, respectively. Stereochemical information was removed, and tautomers were standardized to their canonical forms using the TautomerEnumerator class. Charges were neutralized when possible using a combination of the RDKit Uncharger class and the series of SMARTS patterns provided in the RDKit Cookbook. We found that this approach sometimes incorrectly neutralized functional groups that were written in charge-separated forms, notably sulfoxides and phosphoryl groups, and therefore implemented additional checks to ensure that these functional groups were correctly neutralized. When neutralization altered the net charge of the molecule, both the charged and uncharged forms of the structure were retained for further analysis. Structures containing radical electrons or atoms with invalid valences were removed. The output of this step was a set of deduplicated and standardized two-dimensional structures from PubChem.

We next developed a pipeline to retrieve candidate structures based on the molecular formula of the ground-truth structure. We began by retrieving compounds whose molecular formulas exactly matched that of the ground-truth compound, but found that this approach sometimes failed to retrieve the ground-truth structure or other plausible isomers; moreover, when the same structure was represented in Spectraverse in two different charge states (for instance, as the [M+H]^+^ adduct of a zwitterion versus the [M]^+^ adduct of a compound with a net charge of +1), two different candidate sets would be created. We therefore constructed formula-based candidate sets as follows. First, we extracted the first 14-character block of the InChIKey for the ground-truth compound, and then retrieved from the preprocessed Pub-Chem dataset all entries sharing the same first InChIKey block (representing alternative charge states of this ground-truth structure) with a net charge between –1 and +1 (**Supplementary Fig. 6a**). Manual review of several hundred structures established that the presence of two or more charged moieties that could not be neutralized by the code described above represented structures that were unlikely to occur in nature (**Supplementary Fig. 6b**). We then calculated the molecular formula for each of the structures retrieved by this search, representing valid protonated or deprotonated formulas of the ground-truth structure, and queried PubChem to retrieve all structures corresponding to any of these formulas. If this approach failed to retrieve any candidate structures, PubChem was instead queried with the neutral molecular formula of the ground-truth SMILES, which was adjusted for positively or negatively charged compounds by removing or adding one proton, respectively. The retrieved candidate structures were then deduplicated by grouping structures by the first 14 characters of their InChIKeys and, within each group, retaining the structure with the charge closest to zero (**Supplementary Fig. 6a**). We then ensured that each candidate set contained a structure matching the first InChIKey block of the ground-truth structure, and inserted the ground-truth structure into the candidate set if it was absent. In a small number of cases, a candidate with the same first InChIKey block as the ground-truth structure was present in the candidate set, but with a different canonical SMILES. This occurred when the structure in question was found in PubChem, but only in a different charge state that could not be reached by the series of neutralization steps described above (**Supplementary Fig. 6c**). In these cases, we computed the Tanimoto coefficient to the ground-truth SMILES, and if the similarity was less than 1, replaced the candidate structure with the ground-truth SMILES. Finally, all SMILES with non-zero net charges were passed through the same chemical structure standardization routine described above, and if the resulting SMILES was already present in the candidate set, the charged form was removed. This final step was required to account for a small number of cases in which the charged form of a particular candidate differed in the first InChIKey block from the neutral form, meaning the two structures would not have been identified as duplicates from their InChIKeys alone (**Supplementary Fig. 6d**).

A similar pipeline was developed to generate candidate sets based on the exact mass of the ground-truth structure (simulating experimental scenarios in which the formula of an unidentified small molecule cannot be unambiguously determined). First, the neutral monoisotopic mass for each ground-truth structure was calculated from the precursor m/z and adduct type. We then retrieved from the preprocessed PubChem database all structures whose neutral monoisotopic masses fell within *±*10 ppm of the target mass (after subtracting or adding the mass of a proton for positively or negatively charged compounds, respectively). The resulting candidate sets were then processed as described above for the formula-based candidates. All experiments described in the manuscript used version 1.0.2 of Spectraverse (“Data availability”).

### Generated candidate sets

To generate chemically realistic candidate structures, we trained a chemical language model based on the structured state space sequence (S4) architecture, using the implementation provided by Grisoni and colleagues at https://github.com/molML/s4-for-de-novo-drug-design^67^. Unlike recurrent architectures such as LSTMs, which process sequence elements sequentially, S4 models process the entire sequence simultaneously *via* global convolution during training, an aspect of these models that is purported to help them capture long-range dependencies. During generation, S4 switches to an equivalent recurrent formulation using the same parameters and thereby samples SMILES strings token by token. The model was trained on all ground-truth structures in Spectraverse, which were augmented by enumerating 50 non-canonical SMILES for each structure, using a cross-entropy loss. A total of 100 million SMILES strings were sampled from the trained model, which were then post-processed and deduplicated using the same steps described above for the PubChem-derived candidates. Mass- and formula-based candidate sets were then constructed for the generated structures, following the procedures described above. Chemical similarity between ground-truth and candidate structures was quantified using Tanimoto coefficient on 2048-bit Morgan fingerprints.

We then evaluated the accuracy of structure-only ChemProp models trained to differentiate ground-truth versus candidate structures. We evaluated ChemProp models under both a conventional structure-disjoint data split and when partitioning the data into cross-validation folds based on the MCES distance, but found that only the latter abolished the performance of the structure-only model. To establish the MCES split, we computed the complete matrix of MCES distances between all 44,237 ground-truth structures, and clustered the resulting matrix using single-linkage hierarchical clustering with a minimum MCES distance of 3. Clusters were then aggregated into ten folds of roughly equivalent size using a greedy algorithm. We selected a MCES distance of 3 because it was the highest distance for which the largest cluster produced by single-linkage clustering comprised approximately 10% of the ground-truth structures; more stringent thresholds produced splits in which one fold was considerably larger than all others. The top-*k* accuracy was calculated by ranking candidates in descending order by predicted ground-truth class probabilities, as described above for the MassSpecGym analysis. We also experimented with generating candidates using a LSTM, as previously described and using the implementation provided in https://github.com/skinniderlab/CLM, but found this produced inferior performance (i.e., a structure-only model could more readily distinguish ground-truth structures from candidates generated by the LSTM than by the S4 model; **Supplementary Fig. 7**).

### Benchmarking computational methods for MS/MS annotation

We evaluated the performance of 17 published models for MS/MS annotation on Spectraverse using ten-fold cross-validation on the MCES-disjoint split described above. For each test fold, models were trained on eight folds, with one MCES-disjoint fold reserved as a validation set, and the trained model was then used to rank the candidate structures associated with every spectrum in the test fold. Candidates that were not assigned a score were assigned a similarity of 0. When models assigned identical scores to two or more candidate structures, ties were broken by random permutation to ensure that the ordering of candidates did not influence the results. The Tanimoto coefficient between the ground-truth structure and the top-ranked candidate (top-1 Tc) was calculated using 2048-bit Morgan fingerprints; plots show the top-1 Tc for a random subset of 10,000 spectra. Pairwise comparisons of model performance were performed using paired t-tests across the ten cross-validation folds to test the null hypothesis of no difference in mean top-1 accuracy. Results from formula-based candidate sets are shown in the main text because some models assume the ground-truth formula is known; results from mass-based candidate sets are shown in **Supplementary Fig. 5** but should be interpreted with more caution because they afford an unfair advantage to models that assume a known formula.

#### Collision energy and instrument type

Some MS/MS spectra in Spectraverse were acquired using ramped or stepped collision energies. Consequently, each spectrum is associated with up to three distinct collision energy meta-data fields (typically as a range from minimum to maximum collision energy for ramped acquisitions, and as up to three discrete values for stepped acquisitions). For models that took collision energy metadata as input, we used the mean of these values, expressed either as absolute or normalized collision energy depending on the model. Spectra lacking any reported collision energy were assigned a placeholder value of –1. Similarly, missing instrument types were replaced with the placeholder value “unknown.”

#### MassSpecGym baselines^34^

We evaluated two baseline models proposed by the authors of MassSpecGym, using the implementations provided at https://github.com/pluskal-lab/MassSpecGym. “Fingerprint FFN” is a feed-forward network that accepts a spectrum binned at 1 Da resolution as input and predicts a 4096-bit Morgan fingerprint (radius 2) as output. “DeepSets” processes spectra as sets of raw 2D (m/z and intensity) peak representations and predicts a 4096-bit Morgan fingerprint with radius 2. Both models were trained for 50 epochs, and candidate structures were ranked based on the cosine similarity between predicted and computed fingerprints.

#### ms-pred baselines^18^

We also evaluated two baseline models implemented by Coley and colleagues (using the implementations provided at https://github.com/coleygroup/ms-pred/tree/iceberg_analychem_2024, after modifying them to accommodate additional adducts in the negative ion mode). “Binned spectrum FFN” and “Binned spectrum GNN” (originally “FFN Spec” and “GNN Spec,” renamed here for clarity) are spectrum prediction models that encode input structures using either Morgan fingerprints or a graph neural network, and predict binned spectra at 0.1 Da resolution. Both models were trained for up to 200 epochs, with early stopping triggered if the validation loss failed to improve for 20 and 10 epochs respectively, and candidate structures were ranked based on the cosine similarity between predicted and computed spectra.

#### ChemEmbed^30^

We retrained ChemEmbed models using the implementation provided at https://github.com/massspecdl/ChemEmbed. We implemented the three different versions of ChemEmbed described by Yanes and colleagues: one in which ChemEmbed was trained on individual spectra; one in which ChemEmbed was trained on merged spectra, which were created by combining spectra acquired from the same compound (as identified by the first block of the InChIKey) at multiple collision energies; and a variant of the latter approach in which neutral losses were additionally calculated for each fragment in the merged spectrum. All three were tested on the same set of held-out MS/MS spectra from each fold of Spectraverse. The individual spectrum model performed best, and was used for all results presented in the manuscript. For all three models, spectra were filtered as recommended by the authors of ChemEmbed: fragments more than 0.5 Da above the precursor m/z were removed, as were fragments with intensities less than 1% of the base peak. The remaining peaks were used to construct a binary vector of length 100,000, spanning 0-1,000 Da at 0.01 Da resolution, where value in each bin indicates the presence or absence of a fragment ion at that m/z. All three models were trained to predict a 300-dimensional Mol2vec embedding of the ground-truth structure. Models were trained for up to 70 epochs, with early stopping triggered if the validation loss failed to improve for 10 epochs, and candidate structures were ranked based on the cosine similarity between predicted and computed embeddings

#### ESP^68^

We retrained ESP models using the implementation provided at https://github.com/HassounLab/ESP. Hassoun and colleagues described three versions of ESP: a MLP in which structures are featurized using Morgan fingerprints; a GNN that encodes molecular graphs; and an ensemble model that aggregates predictions made by the MLP and GNN models. The MLP model performed best, and was used for all results presented in the manuscript. This model takes as input a 4096-bit Morgan fingerprint with radius 2, a one-hot vector denoting the adduct type, and the normalized collision energy, and predicts spectra binned at 1 Da resolution. Label mixing and multi-tasking on spectral topic distributions were performed following the default settings in the GitHub repository. Models were trained for 50 epochs, and candidate structures were ranked based on the cosine similarity between predicted and computed spectra.

#### FIORA^29^

We retrained FIORA models using the implementation provided at https://github.com/BAMeScience/fiora. FIORA takes as input a molecular graph as well as metadata (instrument type, adduct, and collision energy in eV); we modified the source code to account for additional adduct types present in Spectraverse, but not the dataset on which FIORA was originally trained. Spectra with missing collision energies were not excluded from the training set. Models were trained for up to 200 epochs, with early stopping triggered if the validation loss failed to improve for 8 epochs, and candidate structures were ranked based on the cosine similarity between predicted and computed spectra after square root transformation of peak intensities.

#### FLARE^31^

We retrained FLARE models using the implementation provided at https://huggingface.co/spaces/HassounLab/FLARE without modifications and following the preprocessing pipeline described in the original implementation (for example, retaining only the 60 most intense fragments and assigning formulas to each fragment using the enumeration-based algorithm from MIST). Models were trained for up to 2,000 epochs, with early stopping triggered if the validation loss failed to improve for 300 epochs, and candidate structures were ranked based on the bidirectional average of peak-to-atom and atom-to-peak similarities.

#### FraGNNet^14^

We retrained FraGNNet models using the version 1.5 implementation provided at https://github.com/FraGNNet/fragnnet/tree/release/v1.5. This implementation differs from that described by Young et al.^14^ in two respects: (1) the model takes as input additional metadata, including instrument type and fragmentation mode (HCD, CID, or unknown), and (2) the model explicitly models charge-migration fragmentation to better support adduct types such as [M+Na]^+^ and [M+Cl]^−69^. As in the MassSpecGym benchmark^34^, we use the computationally efficient D3 NodeMLP variant rather than the original D4 model, which reduces the size of the generated DAG (and therefore the cost of spectrum prediction) at the expense of accuracy. Models were trained for up to 100 epochs, and the checkpoint with the lowest validation loss was selected. Candidate structures were ranked based on the Jensen-Shannon similarity between predicted and computed spectra after square root transformation.

#### GrAFF-MS^16^

We retrained GrAFF-MS models using the implementation provided at https://github.com/coleygroup/ms-pred/tree/iceberg_analychem_2024 without modifications. Models were trained for up to 200 epochs, with early stopping triggered if the validation loss failed to improve for 20 epochs, and candidate structures were ranked based on the cosine similarity between predicted and computed spectra.

#### ICEBERG^17^

We retrained ICEBERG models using the implementation provided at https://github.com/coleygroup/ms-pred/tree/iceberg_analychem_2024 (i.e., the implementation reported by ref.^17^) after modifying the code to accommodate additional adducts in the negative ion mode. Models were trained for up to 200 epochs, with early stopping triggered if the validation loss failed to improve for 20 epochs, and candidate structures were ranked based on the cosine similarity between predicted and computed spectra.

#### JESTR^23^

We retrained JESTR models using the implementation provided at https://github.com/HassounLab/JESTR1 and following the preprocessing pipeline adopted in this repository, but omitting the regularization loss because the degree and timing regularization was carefully selected in the original publication (10% weight given to the regularization loss for the last 3% of the training epochs only), and would likely require careful tuning in Spectra-verse (e.g., using a nested cross-validation approach). Models were trained for up to 1,000 epochs, with early stopping triggered if the validation loss failed to improve for 80 epochs, and candidate structures were ranked based on the cosine similarity between the embeddings of query spectra and candidate structures.

#### MassFormer^15^

We retrained MassFormer models using the implementation provided at https://github.com/coleygroup/ms-pred/tree/iceberg_analychem_2024 after modifying the code to accommodate additional adducts in the negative ion mode. Spectra were converted into fixed-length vectors by binning at 0.1 Da resolution, up to a maximum m/z of 1,500 Da. Models were trained for up to 200 epochs, with early stopping triggered if the validation loss failed to improve for 20 epochs, and candidate structures were ranked based on the cosine similarity between predicted and experimental MS/MS spectra.

#### MetFID^27^

We reimplemented MetFID based on the description provided by Ressom and colleagues, for which no code was released. Preprocessing followed the steps described in the MetFID publication: fragment intensities were scaled such that all intensity values were between 0 and 100; fragments with m/z values exceeding the precursor m/z were identified and any other fragments whose intensity was less than the maximum of these ions were removed; other fragments with relative intensities less than 10% were removed; spectra for each compound (as identified by the first InChIKey block) were merged to obtain a single consensus spectrum for compound, grouping peaks with a 0.1 Da m/z tolerance and averaging their intensities; and neutral loss features were derived from the consensus spectra, using the intensities of each fragment ion as a proxy for the corresponding neutral loss. Spectra with fewer than five peaks reaching at least 2% relative intensity were discarded as uninformative, as recommended by the authors. Both the fragment ion peaks and neutral-loss features were independently binned at a resolution of 1 Da over the range 0-1000 m/z and were concatenated into a single input feature vector. Bins with zero intensity across the entire training set were removed. Fingerprints were calculated for each compound in the training set using a concatenation of MACCS keys (computed via RDKit), FP3, and FP4 fingerprints (the latter two both computed via OpenBabel). An artificial neural network with two hidden layers of sizes 800 and 600 was trained to predict this fingerprint, minimizing the cross-entropy loss. Models were trained for 30 epochs, and candidate structures were ranked based on the mean absolute error (MAE) between computed and predicted fingerprints.

#### MIST^20^

We retrained MIST models using the implementation provided at https://github.com/samgoldman97/mist, after modifying the code to accommodate additional adducts in the negative ion mode. No data augmentation with predicted spectra was performed. We evaluated two retrieval strategies for MIST: ranking candidates directly by their predicted fingerprints, or ranking them using a learned distance metric from a final contrastive fine-tuning step. The latter approach yielded better performance, so was used for all results reported here. Models were trained for up to 600 epochs, with early stopping triggered if the validation loss failed to improve for 20 epochs, and candidate structures were ranked based on the cosine similarity between the embeddings of query spectra and candidate structures.

#### NEIMS^26^

We retrained NEIMS models using our own reimplementation that follows the architecture of the original model. The model consists of a four-layer residual MLP that passes 4096-bit Morgan fingerprints with radius 2 through a bidirectional spectrum FFN with a 0.5 bottleneck factor to predict binned spectra at 1 Da resolution. Because no mechanism for conditioning on adduct type was described in the original publication, we trained independent NEIMS models for each adduct type in Spectraverse. Models were trained for up to 50 epochs, and the checkpoint with the lowest validation loss was selected. Candidate structures were ranked based on the cosine similarity between predicted and computed spectra.

#### 3DMolMS^28^

We retrained 3DMolMS models using the implementation provided at https://github.com/coleygroup/ms-pred/tree/iceberg_analychem_2024 after modifying the code to accommodate additional adducts in the negative ion mode. Spectra were converted into fixed-length vectors by binning at 0.1 Da resolution, up to a maximum m/z of 1,500 Da. Models were trained for up to 200 epochs, with early stopping triggered if the validation loss failed to improve for 20 epochs, and candidate structures were ranked based on the cosine similarity between predicted and experimental MS/MS spectra

#### Nearest-neighbor baselines

To further place the performance of published machine-learning models in context, we implemented a nearest-neighbor baseline recently proposed by Khoo and Barzilay^60^. In this approach, each spectrum in the test fold is assigned the chemical fingerprint associated with its nearest-neighbor in the training folds (using 4096-bit Morgan fingerprints with radius 2). Whereas Khoo and Barzilay evaluated the accuracy of this predicted fingerprint directly using the Jaccard index, we instead ranked the candidate structures for each spectrum based on their cosine similarity between their computed fingerprints and this nearest-neighbor fingerprint, so as to evaluate this baseline in the setting of the retrieval task evaluated here (recognizing that retrieval of the ground-truth structure, which is the task ultimately of interest, may not be correlated and may in fact be anticorrelated with the accuracy of fingerprint prediction^70^). Moreover, we extend this baseline by identifying nearest neighbors using three measures of spectral similarity (cosine, neutral loss cosine, and modified cosine similarity), using the implementations from https://github.com/YuanyueLi/SpectralEntropy (cosine similarity)^71^ and matchms (neutral loss and modified cosine similarity)^72^, respectively, and report the performance of each variant separately. For the cosine similarity, spectra in the training folds were not filtered based on the precursor m/z of the test spectrum (i.e., we performed an open cosine search).

#### Rank product aggregation

To combine the predictions of multiple methods into a single consensus ranking, we adapted the rank-product approach originally introduced for meta-analysis of transcriptomic experiments^73^. This is a deliberately simple approach that was selected in order to probe whether different methods provide complementary information (and not as an attempt to identify an optimal aggregation strategy, which is left for future work). For a given query spectrum, each method independently assigns a rank to each candidate structure. We then aggregated these predictions by computing the product of per-method ranks for each candidate structure, considering varying numbers of top-scoring methods (as determined by their top-1 accuracy). Candidates were then re-ranked in ascending order of their rank product to yield the consensus ranking, such that candidates ranked highly by many methods were prioritized. Ties were broken at random.

## Data availability

Spectraverse v1.0.2, including MS/MS spectra, their associated metadata, and all language model-generated candidate sets, is available from Zenodo at https://doi.org/10.5281/zenodo.19927403. Results from our benchmark of 17 methods, including the scores assigned to each structure in candidate sets for held-out spectra, are available from Zenodo at https://doi.org/10.5281/zenodo.21089924.

---

**Supplementary Fig. 1.**
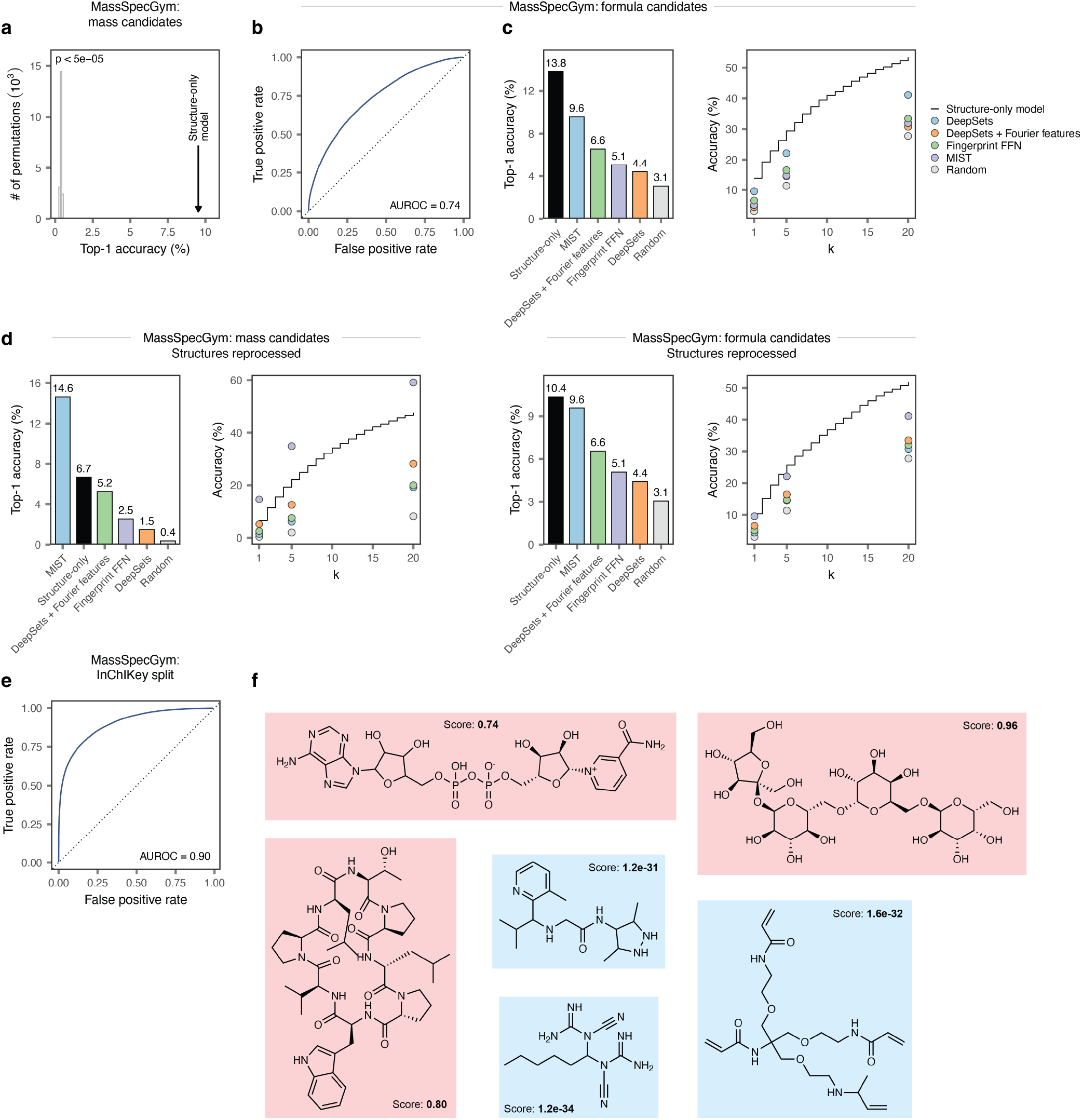
Supporting data for the performance of the structure-only model in MassSpecGym. **a**, Statistical significance of the performance of the structure-only model, as compared to a random baseline (20,000 bootstraps). **b**, Receiver operating characteristic (ROC) curve showing the performance of the structure-only model in distinguishing between ground-truth and candidate structures in the held-out test fold of MassSpecGym, as in **Fig. 1c** but here for the bonus challenge of MassSpecGym, in which the molecular formula is known. Inset text shows the area under the ROC curve (AUROC). **c**, Top-1 accuracy, left, or top-*k* accuracy (for *k* ≤ 20), right, of the structure-only model, as compared to baselines reported in MassSpecGym, as in **Fig. 1d** but here for the bonus challenge of MassSpecGym in which the molecular formula is known. **d**, As in **c**, but after reprocessing all ground-truth and candidate structures to ensure uniform molecule standardization and handling of tautomers. **e**, Receiver operating characteristic (ROC) curve showing the performance of the structure-only model in distinguishing between ground-truth and candidate structures, when splitting spectra and candidate sets in MassSpecGym by MCES versus a more conventional structure-disjoint split. **f**, Additional examples of test set compounds predicted to be ground-truth or candidate structures with high confidence by the structure-only model.

**Supplementary Fig. 2.**
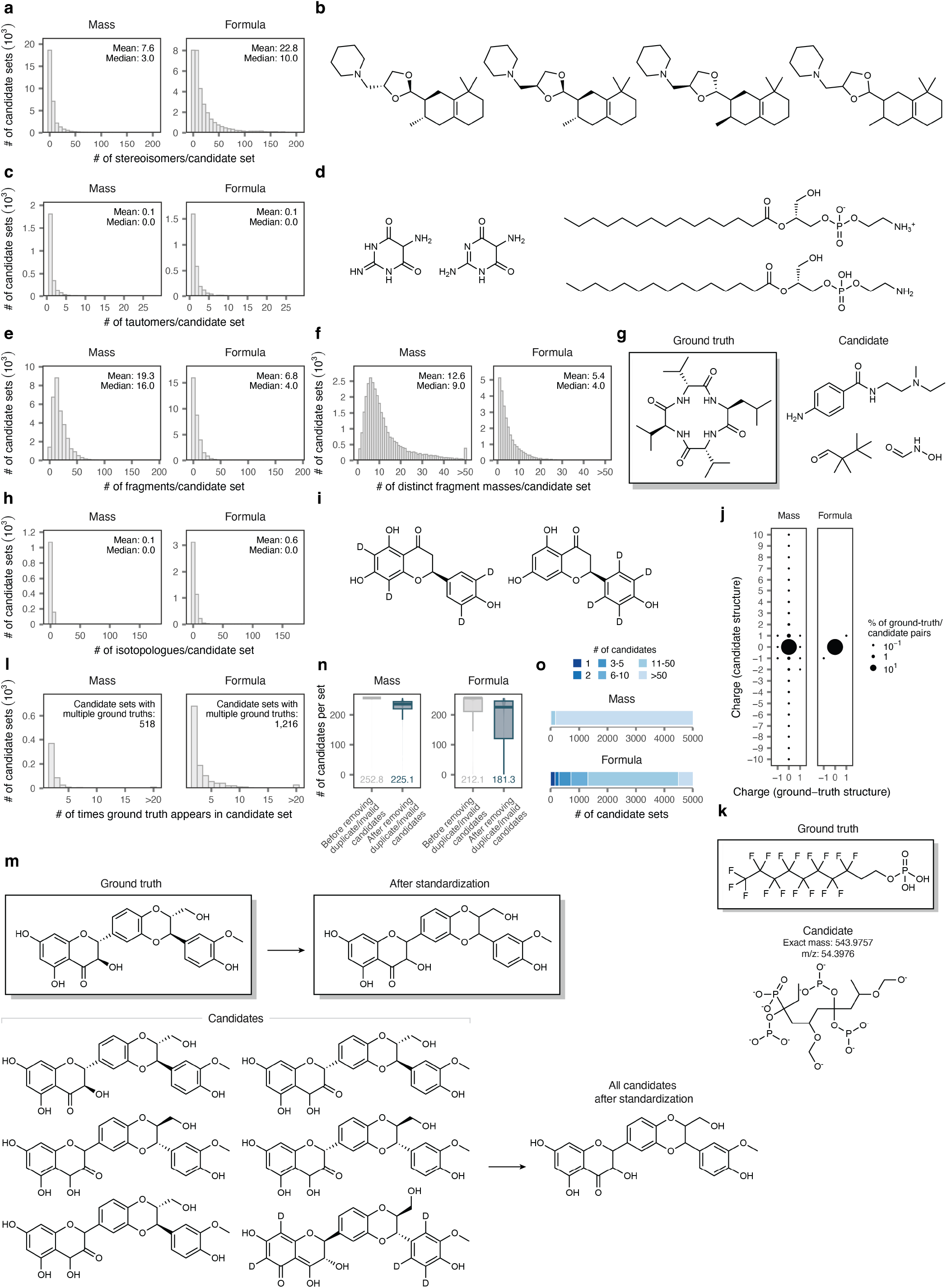
MassSpecGym candidate sets. **a**, Number of structures per candidate set that are stereoisomers of another candidate structure, shown separately for mass or formula candidates. **b**, Example of four stereoisomers present in the same candidate set. **c**, Number of structures per candidate set that are tautomers of another candidate structure, shown separately for mass or formula candidates. Candidate sets containing zero tautomers are removed to improve visualization. **d**, Two examples of tautomeric structures present in the same candidate set. **e**, Number of structures per candidate set that are composed of multiple disconnected fragments, shown separately for mass or formula candidates. **f**, Number of unique masses per candidate set (after rounding to two decimal places) when retaining only the largest fragment within each structure. **g**, Example of a ground-truth and candidate structure pair in which the candidate structure is composed of multiple disconnected fragments. **h**, Number of structures per candidate set that are isotopologues of another candidate structure, shown separately for mass or formula candidates. Candidate sets containing zero isotopologues are removed to improve visualization. **i**, Example of two isotopomeric structures present in the same candidate set. **j**, Comparison of net charges for ground-truth versus candidate structures, shown separately for mass or formula candidates. Charges greater than +10 or less than –10 are winsorized to +10 and –10, respectively. **k**, Example of a ground-truth and candidate structure pair in which the candidate structure is implausible due to its charge state. **l**, Number of candidate sets containing two or more redundant representations of the ground-truth structure. In these candidate sets, models would thus be arbitrarily penalized or rewarded for favoring one representation over the other. **m**, Example of a candidate set in which the ground-truth structure appears six times as various different stereoisomers, tautomers, or isotopologues. **n**, Number of structures per candidate set, before and after removal of duplicate and implausible structures, shown separately for mass or formula candidates. **o**, Number of candidate sets containing less than 50 structures, shown separately for mass or formula candidates.

**Supplementary Fig. 3.**
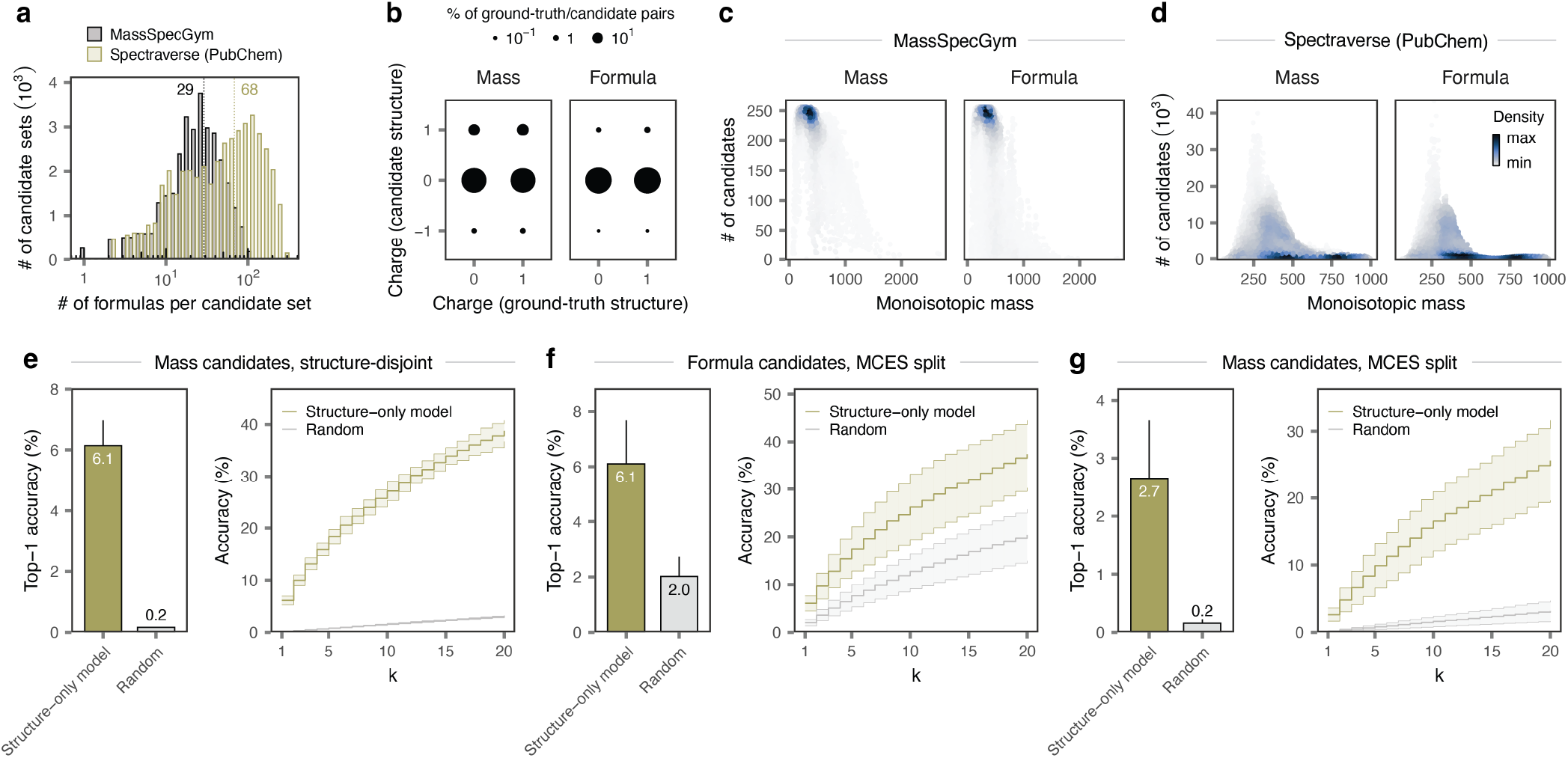
Supporting data for expanded PubChem candidate sets. **a**, Number of distinct molecular formulas within each unique mass-based candidate set in MassSpecGym versus the expanded candidate sets retrieved from PubChem for MS/MS spectra in Spectraverse. **b**, Comparison of net charges for ground-truth versus candidate structures, for PubChem-derived candidate sets. **c**, Relationship between the size of each candidate set and the monoisotopic mass of the ground-truth structure in MassSpecGym. **d**, As in **c**, but for expanded candidate sets retrieved from PubChem for MS/MS spectra in Spectraverse. **e**, Top-1 accuracy, left, or top-*k* accuracy (for *k* ≤ 20), right, of the structure-only model in expanded PubChem candidate sets, as compared to a random selection baseline, for mass-based candidate sets in structure-disjoint cross-validation. Error bars show standard deviation across ten cross-validation folds. **f**, As in **e**, but for formula-based candidate sets with a MCES distance split. **g**, As in **e**, but for mass-based candidate sets with a MCES distance split.

**Supplementary Fig. 4.**
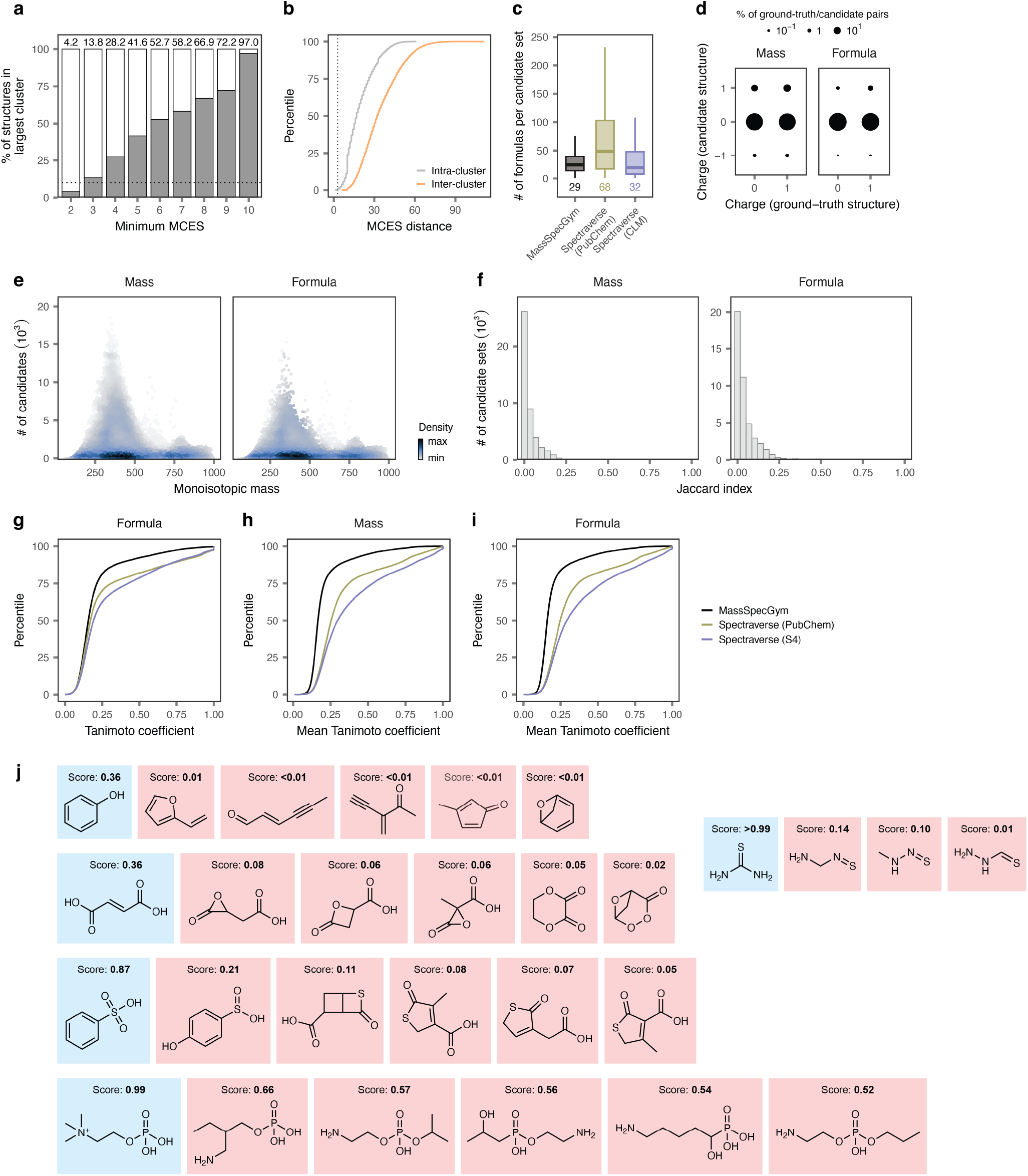
Supporting data for language model-generated candidate sets. **a**, Proportion of ground-truth structures in the single largest cluster, when clustering the matrix of MCES distances between all ground-truth structures using single-linkage hierarchical clustering across a range of MCES distance thresholds. **b**, Distribution of intra-vs inter-cluster MCES distances in the final MCES split. Dotted line shows a MCES distance of 3. **c**, Number of distinct molecular formulas within each unique mass-based candidate set in MassSpecGym versus PubChem-derived and language model-generated candidate sets for MS/MS spectra in Spectraverse. **d**, Comparison of net charges for ground-truth versus candidate structures, for language model-generated candidate sets. **e**, Relationship between the size of each language model-generated candidate set and the monoisotopic mass of the ground-truth structure in Spectraverse. **f**, Distribution of Jaccard indices between PubChem-derived and language model-generated candidate sets for MS/MS spectra in Spectraverse. **g**, Cumulative distribution function of Tanimoto coefficients between ground-truth structures and one randomly selected structure per candidate set, for MassSpecGym, PubChem-derived candidate sets, or language model-generated formula-based candidate sets. **h**, As in **g**, but showing the mean Tanimoto coefficient of the top-255 nearest neighbors for mass-based candidate sets. **i**, As in **g**, but showing the mean Tanimoto coefficient of the top-255 nearest neighbors for formula-based candidate sets. **j**, Additional examples of generated candidate sets in which the ground-truth compound can be deduced by the structure-only model without access to MS/MS information.

**Supplementary Fig. 5.**
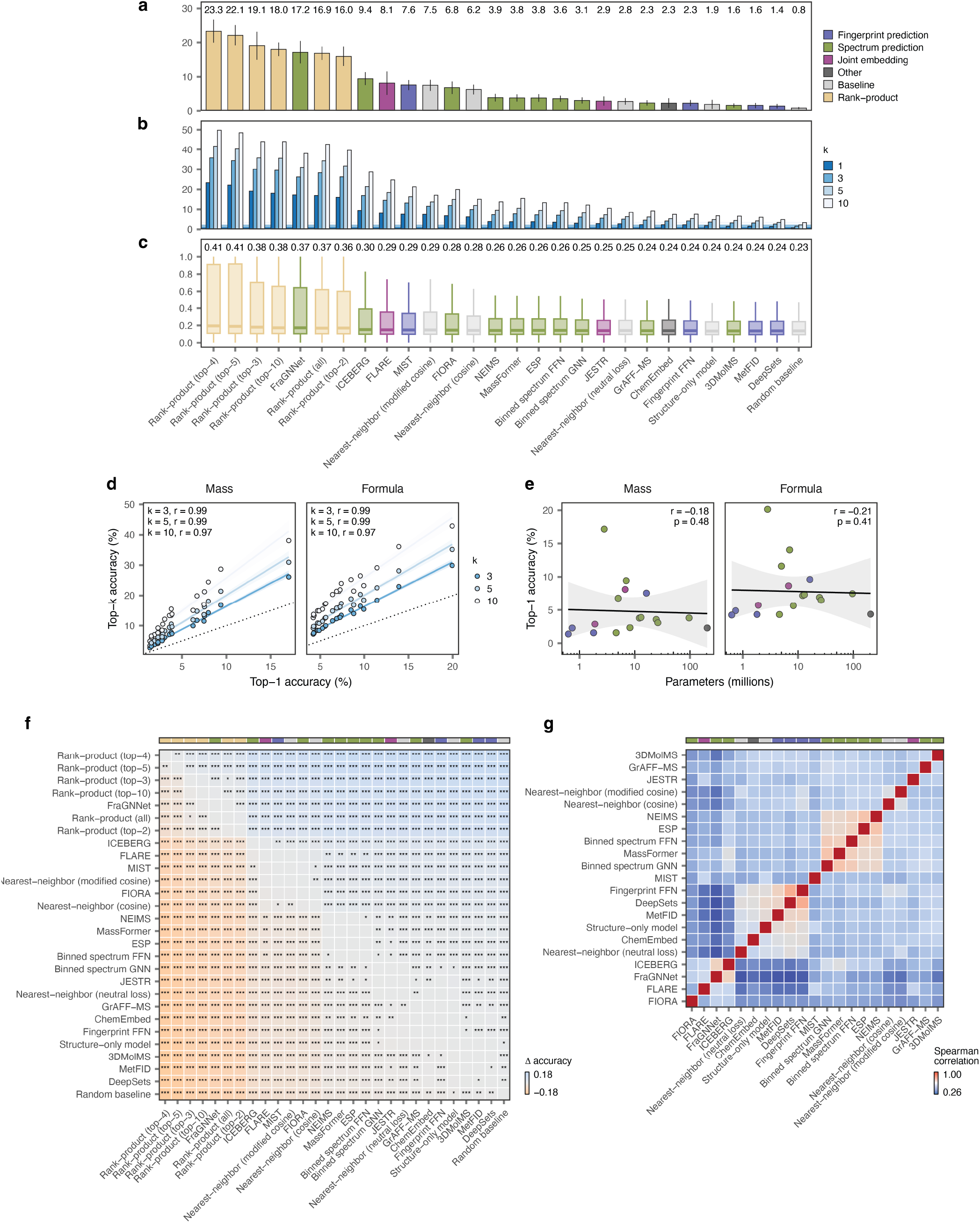
Additional benchmarking of computational methods for MS/MS annotation. **a**, Top-1 accuracy of 17 published models for MS/MS annotation, baseline approaches, and rank-product aggregation of varying numbers of top-performing models, as in **Fig. 4a** but here showing mass-based candidate sets. Error bars show standard deviation across ten cross-validation folds. Inset text shows the mean top-1 accuracy across folds. **b**, As in **a**, but showing the top-3, top-5, and top-10 accuracy for each model. **c**, As in **a**, but showing the Tanimoto coefficient between the ground-truth structure and the top-ranked candidate. Inset text shows the mean Tanimoto coefficient. **d**, Correlation between top-1 and top-3, top-5, or top-10 accuracy for the models and baselines shown in **a**. **e**, Relationship between top-1 accuracy and parameter count for the 17 published models for MS/MS annotation shown in **a**. **f**, Mean difference in the top-1 accuracy (Δ accuracy) between the models shown in **a**. Text highlights comparisons with a two-sided paired t-test p-value less than 0.05 (*, p < 0.05; **, p < 0.01; ***, p < 0.001). **g**, Spearman correlation matrix between ranks assigned to ground-truth structures by the 17 published models for MS/MS annotation shown in **a**.

**Supplementary Fig. 6.**
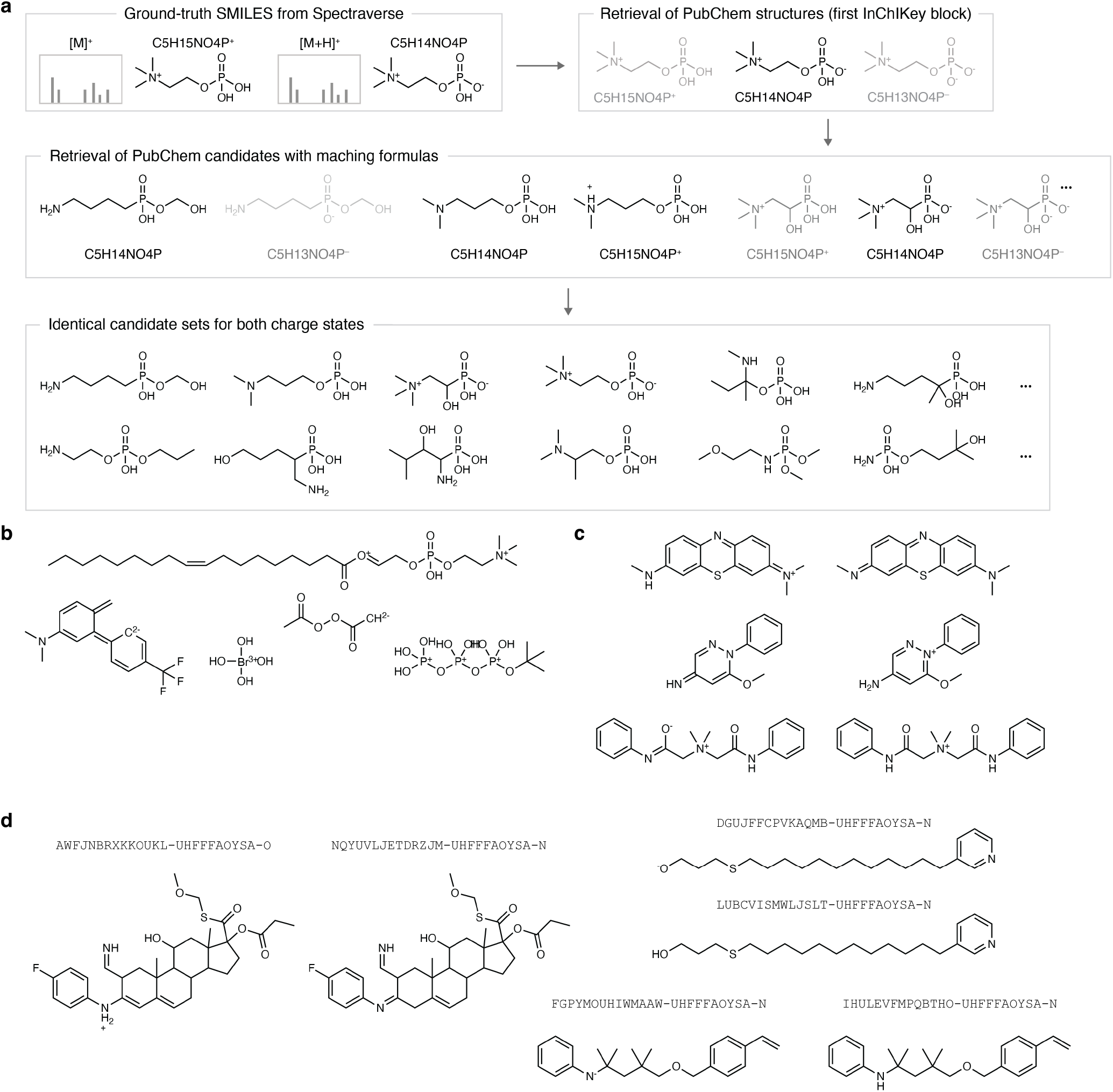
Creation of PubChem candidate sets for MS/MS spectra in Spectraverse. **a**, Overview of formula-based candidate set creation, accounting for the charge state of the ground-truth compound such that compounds represented in different charge states in Spectraverse have identical candidate sets. **b**, Examples of candidate structures with absolute net charges of two or greater that could not be neutralized by our approach (and were therefore discarded). **c**, Examples of cases in which a candidate with the same first InChIKey block as the ground-truth structure was present in the candidate set, but with a different canonical SMILES (and a Tanimoto coefficient less than 1). **d**, Examples of candidate structures for which the charged form differed in the first InChIKey block from the neutral form.

**Supplementary Fig. 7.**
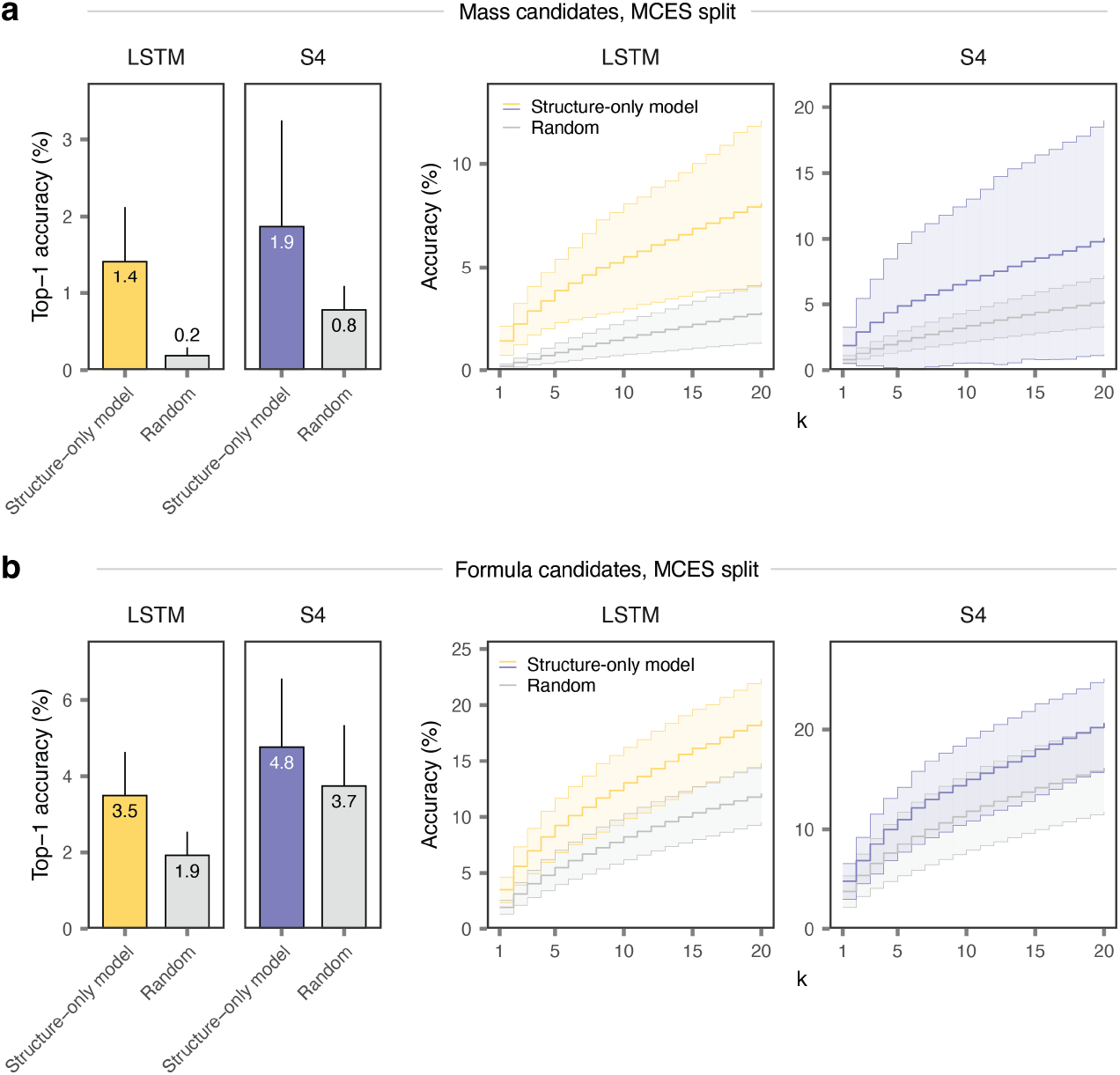
Benchmarking language model architectures for the generation of candidate structures. **a**, Top-1 accuracy, left, or top-*k* accuracy (for *k* ≤ 20), right, of the structure-only model in candidate sets generated by language models based on a LSTM versus S4 architecture, as compared to a random selection baseline, for mass-based candidate sets with a MCES distance split. Error bars and shaded areas show standard deviation across ten cross-validation folds. **b**, As in **a**, but for formula-based candidate sets.

